# Prolyl hydroxylation is required for maturation and secretion of Zika virus non-structural protein 1

**DOI:** 10.64898/2026.02.12.705510

**Authors:** Bodine Bezemer, Gijs J. Overheul, Milou Stevens, Cornelia G. Spruijt, Michiel Vermeulen, Pascal Miesen, Ronald P. van Rij

## Abstract

Mosquito-borne orthoflaviviruses such as Zika virus (ZIKV) and dengue virus (DENV) have a high epidemic potential and are associated with a wide range of possible outcomes, from asymptomatic infections to severe complications. The orthoflavivirus non-structural protein 1 (NS1), which can be membrane-bound as well as secreted from the cell, plays important roles in viral replication, immune evasion and pathogenesis. To describe the interactome of ZIKV NS1, we infected Huh7 and neural progenitor cells with a recombinant ZIKV expressing FLAG-tagged NS1 and identified interacting proteins by quantitative mass spectrometry. In both cell types, multiple prolyl hydroxylases, glycosylation enzymes, ER-associated degradation proteins, and proteins involved in immunity were identified as NS1 interactors. Mass spectrometry data showed that NS1 contains potential hydroxyproline residues, suggesting that prolyl hydroxylase enzymes are involved in post-translational modification of NS1. Chemical inhibition of prolyl hydroxylases impaired NS1 intracellular trafficking and plasma membrane localization and reduced ZIKV RNA levels and titers. Mutagenesis of the proline residue at position 281 impaired NS1 secretion, although viral replication was unaffected. Moreover, substituting proline at position 267 strongly dysregulated NS1 expression and reverted to the wild-type sequence already after a single passage, suggesting that this mutation leads to severe replication defects. Our results identify prolyl hydroxylation as an essential post-translational modification of ZIKV NS1.

## Introduction

The mosquito-borne Zika virus (ZIKV) was long regarded as a relatively unimportant human pathogen, since most reported infections were asymptomatic or only led to mild disease^1^. This perception changed during the outbreaks in the Americas in 2015 and 2016, which were associated with microcephaly and other congenital malformations^1^. ZIKV has a positive-sense, single-stranded RNA genome that codes for a single polyprotein, which is cleaved into the three structural proteins, capsid (C), premembrane (prM) and envelope (E), and the seven non-structural genes NS1, NS2A, NS2B, NS3, NS4A, NS4B and NS5. The structural proteins are necessary for entry and virion assembly and the non-structural proteins are involved in viral RNA replication, polyprotein processing, and host-pathogen interactions^2^.

The NS1 protein plays a complex role in viral infection, pathogenesis and immune evasion^3,4^. Upon translation of the viral polyprotein, NS1 monomers are expressed in the ER lumen where they become glycosylated and dimerize before associating with the ER membrane^5-7^. The ER membrane associated NS1 dimers have an incompletely characterized role in viral RNA replication, but they are proposed to remodel the ER and provide a structural scaffold for replication complex formation^8,9^. These dimers also migrate to the plasma membrane, where they associate with cholesterol and lipid rafts^6,10,11^. In addition, NS1 dimers are transported to the Golgi apparatus, where the glycans are modified into more complex carbohydrates and three dimers can form a channel-like hexamer containing a lipid cargo in the center^3,12^. Following secretion, these hexamers contribute to pathogenesis by inducing the destruction of the endothelial glycocalyx layer and promoting immune evasion by inhibition of the complement system^13,14^.

Identification of binding partners of NS1 may yield insights into the processes underlying proper NS1 folding, transport, and function. However, most orthoflavivirus NS1 interactome studies have relied on the overexpression of NS1 proteins or on subgenomic replicon systems^15-19^, which may not accurately reflect conditions in an infected cell. To identify the host proteins that interact with NS1 in the context of an authentic infection, we generated a ZIKV recombinant containing a single internal FLAG-tag in NS1. Quantitative mass spectrometry analysis of cells infected with the FLAG-tagged virus revealed a network of collagen prolyl hydroxylases as NS1 binding host factors. Collagen prolyl hydroxylases are enzymes that convert proline residues into hydroxyproline to regulate the stability of their protein substrates, such as collagens and other proteins including the RNAi effector protein Argonaute 2^20,21^. Using NS1 mutants and pharmacological inhibition of prolyl hydroxylases, we show that prolyl hydroxylation is required for ZIKV NS1 maturation, and secretion.

## Results

### NS1 interactome in ZIKV infected cells

Orthoflavivirus NS1 is involved in processes ranging from viral replication to pathogenesis and immune evasion^4^, but the molecular mechanisms behind these processes are not fully elucidated. We aimed to identify host factors interacting with ZIKV NS1 in infected cells by quantitative mass spectrometry. We used circular polymerase extension reaction (CPER) cloning to introduce an internal FLAG-tag in NS1 after amino acid position 342 (Fig. 1a), a site found to be permissive to insertions in a transposon mutagenesis screen^22^. To confirm that ZIKV containing FLAG-tagged NS1 (ZIKV^NS1:FLAG^) does not have a major replication defect, we compared its replication kinetics to that of a CPER-generated wildtype ZIKV without an insertion (ZIKV^WT^) in Huh7 cells. Indeed,the replication kinetics of both CPER-generated viruses were similar (Fig. 1b, c), indicating that the FLAG-tag does not significantly impair virus replication and that the ZIKV^NS1:FLAG^ can be used for our study. Viral RNA levels and titers of the CPER-generated viruses were lower than those of the clinical isolate H/PF/2013, which was probably due to the use of another isolate, BeH819015, for the production of the CPER viruses^23^.

**Figure 1.**
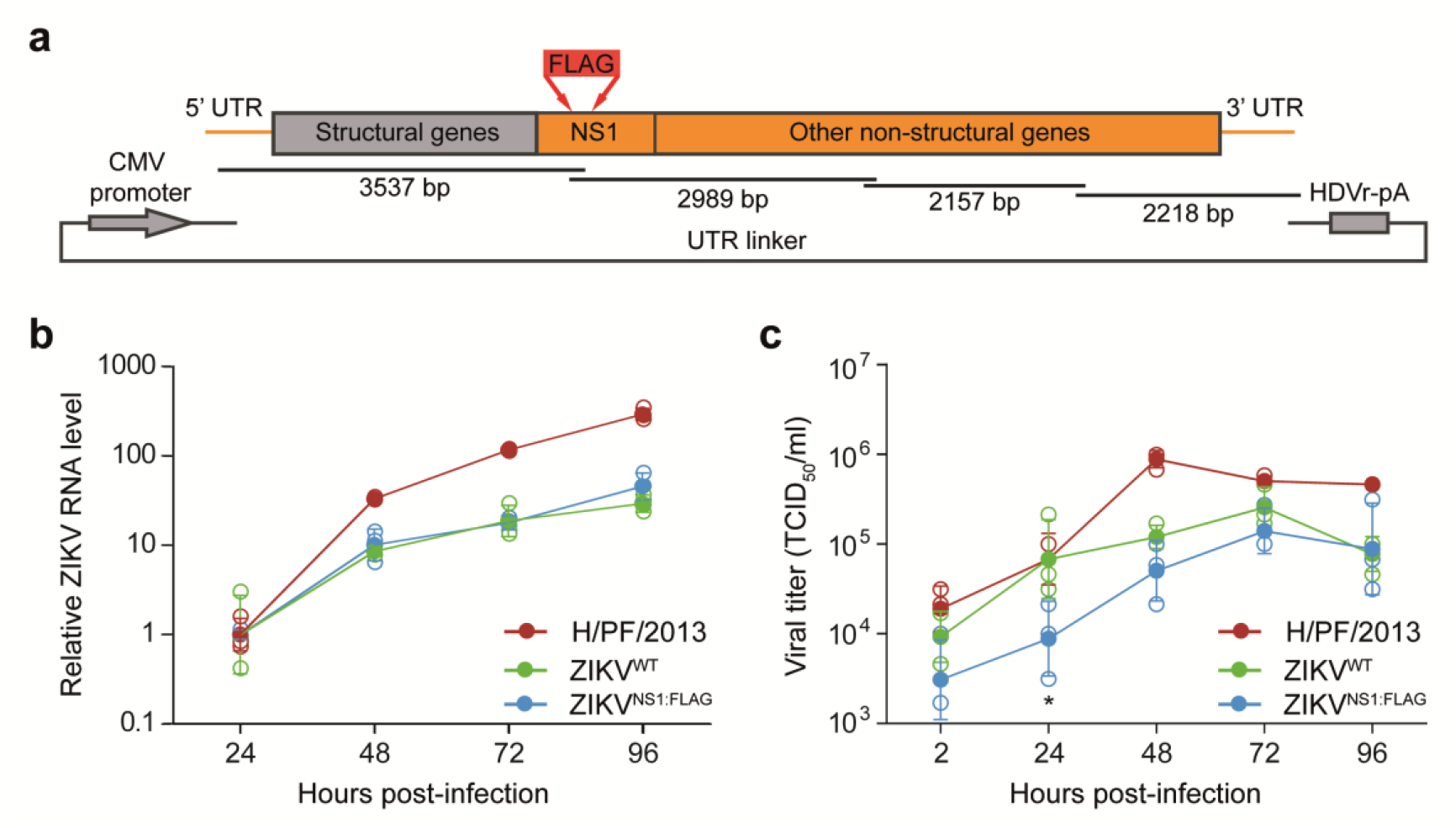
Creation and validation of the CPER-based NS1-tagged ZIKV. **a** Schematic of the circular polymerase extension reaction (CPER) strategy to create ZIKV recombinant virus with a FLAG-tag in NS1. The FLAG-tag between amino acid position 342 and 343 is represented by a red arrow. HDVr-pA, hepatitis delta virus ribozyme followed by a poly(A) signal. **b,c** ZIKV RNA levels (**b**) and infectious titers (**c**) in Huh7 cells at different time points post-infection with ZIKV H/PF/2013, ZIKV^WT^ or ZIKV^NS1:FLAG^ at an MOI of 0.1. ZIKV RNA levels were measured by RT-qPCR (**b**) and normalized to ZIKV RNA levels at 24 hours post-infection for each virus. Titers measured at 2 hours post-infection reflect residual input virus. Data of ZIKV^WT^ and ZIKV^NS1:FLAG^ were log-transformed and analyzed with one-way ANOVA with Holm-Šidák post-hoc test (* *p* < 0.05, other comparisons were not significant). H/PF/2013 was not included in the statistical analysis. Data represent mean (filled symbols) and SD of three biological replicates, open symbols represent individual data points.

We investigated the interactome of NS1 in two cell types, the immortalized human hepatocellular carcinoma Huh7 cell line and human induced pluripotent stem cell-derived neural progenitor cells (NPCs), infection of which results in dysregulated neurogenesis, one of the major causes of ZIKV-induced birth defects^24-27^. We infected the cells for 48 hours with ZIKV^WT^ or ZIKV^NS1:FLAG^ and enriched for NS1 interactors by FLAG immunoprecipitation (IP). At the time of lysis, viral titers were comparable for both viruses in each cell type (Supplementary Fig. 1). Label-free quantitative mass spectrometry of the immunoprecipitated proteins identified 36 significantly enriched proteins in FLAG IP samples of ZIKV^NS1:FLAG^ infected NPCs. As expected, these proteins included the NS1 bait protein as well as the viral E protein, known to interact with flavivirus NS1^28^, validating our approach (Fig. 2a, Supplementary Table 2, Supplementary Table 3). For Huh7 cells, 27 proteins were significantly enriched in the ZIKV^NS1:FLAG^ IP samples, including the NS1 bait protein, the viral E protein as well as prM, previously reported to interact with flavivirus NS1 for viral particle formation^28^ (Fig. 2b, Supplementary Table 2, Supplementary Table 4). In contrast, NS5 was depleted from ZIKV^NS1:FLAG^ IP, in line with the different subcellular localization of these viral proteins^28^.

**Figure 2.**
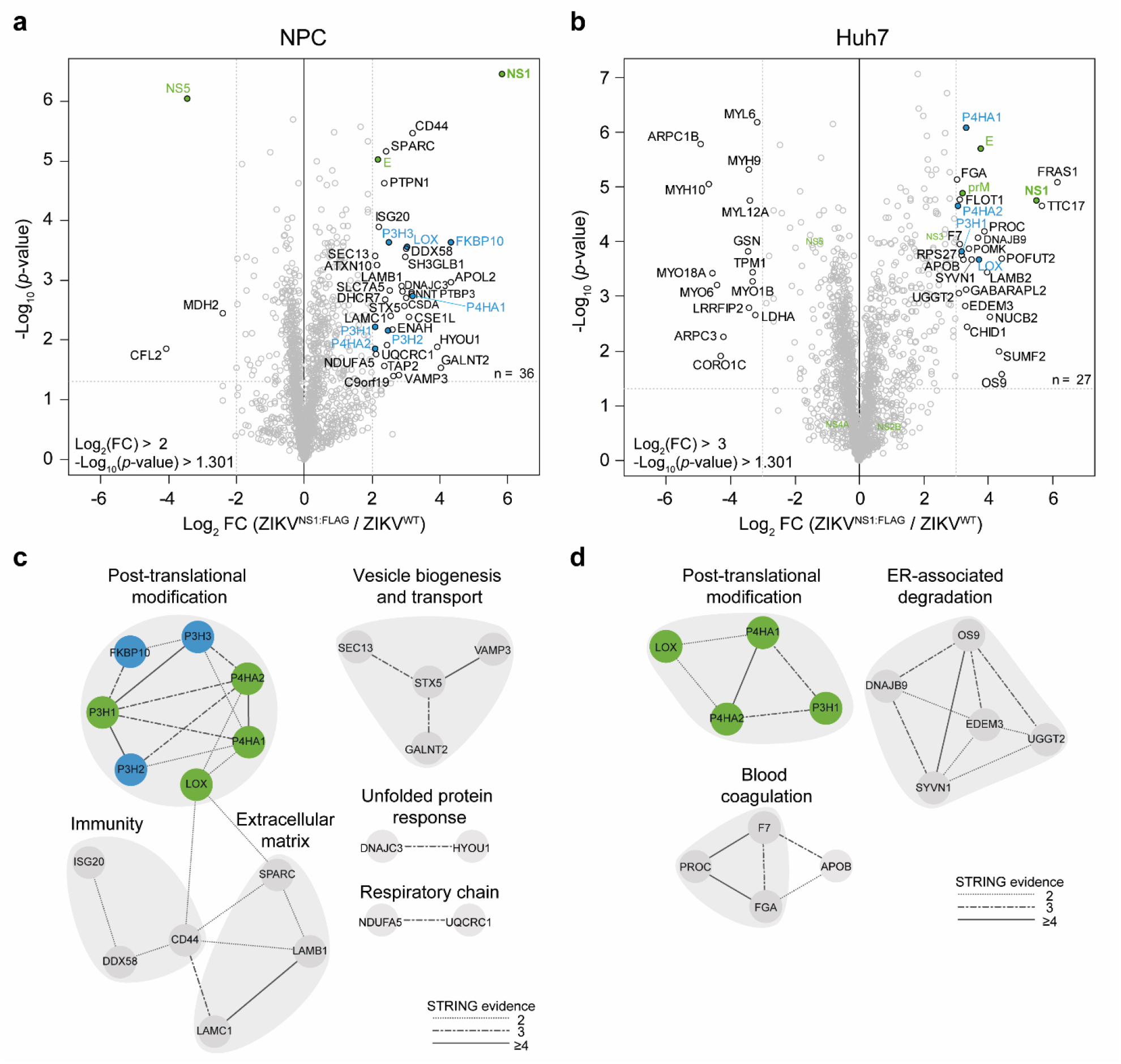
NS1 interacts with prolyl hydroxylases. **a, b** Volcano plots of the NS1 interactors in ZIKV^WT^ or ZIKV^NS1:FLAG^ infected NPCs (a) and Huh7 cells (**b**) after FLAG IP using ZIKV^WT^ infected cells as a control. Interacting proteins were detected by label-free quantitative mass spectrometry. The X-axis shows the log_2_ fold change (FC) in FLAG IP from cells infected with ZIKV^NS1:FLAG^ over cells infected with ZIKV^WT^; the Y-axis shows the -Log10*(p-*value) in two sample *t*-tests. Dashed lines correspond to the cutoff values for the log_2_ FC and the -log_10_ *p*-value, and specific interactors are shown in the top right quadrant. Viral proteins are shown in green and proteins involved in post-translational modification are shown in blue. The indicated number of interactors include the viral proteins. **c, d** STRING analysis of NS1 interacting proteins in NPCs (**c**) or Huh7 cells (**d**). Green nodes in the post-translational modification network represent proteins interacting with NS1 in both cell types and blue nodes represent proteins that were only enriched in NPCs. Edges are stylized according to the number STRING evidence types supporting each interaction (homology, co-expression, experimentally determined interactions, database annotation or automated text mining).

### ZIKV NS1 interacts with collagen prolyl hydroxylases

We performed STRING and gene ontology (GO) term analyses to explore the protein-protein interaction network associating with NS1. In line with the known ER localization of NS1, ER-localized proteins were enriched in NS1 IP both in NPCs (15/33 host proteins) and Huh7 cells (15/24 host proteins) (Supplementary Table 5, 6). Moreover, the GO term endomembrane system, which besides the ER includes the Golgi apparatus and transport vesicles, was associated with 23 and 20 NS1 interactors in NPC and Huh7 cells, respectively, suggesting that we also identified proteins interacting with NS1 during its transport through the Golgi towards the plasma membrane. As expected, we found four enzymes involved in glycosylation (Polypeptide N-acetylgalactosaminyltransferase 2, GALNT2; Protein O-mannose kinase, POMK; GDP-fucose protein O-fucosyltransferase 2, POFUT2; UDP-glucose:glycoprotein glucosyltransferase 2, UGGT2)^5^ as well as three proteins that function in immunity (Antiviral innate immune response receptor RIG-I, DDX58; Antigen peptide transporter 2, TAP2; Interferon-stimulated gene 20 kDa protein, ISG20) in NPCs (Fig. 2c, d). Moreover, in line with previous observations for dengue virus (DENV) NS1^15,18^, we detected five proteins involved in ER-associated degradation in Huh7 cells (UDP-glucose:glycoprotein glucosyltransferase 2, UGGT2; Protein OS-9, OS9; E3 ubiquitin-protein ligase synoviolin, SYVN1; ER degradation-enhancing alpha-mannosidase-like protein 3, EDEM3; DnaJ homolog subfamily B member 9, DNAJB9, Fig. 2d) as well as proteins involved in the unfolded protein response pathway during ER stress in NPCs (DnaJ homolog subfamily C member 3, DNAJC3; Hypoxia up-regulated protein 1, HYOU1). NS1 also associated with extracellular matrix proteins including subunits of the heterotrimeric laminin complex, laminin beta-1 (LAMB1) and laminin gamma-1 (LAMC1) in NPCs (Fig. 2c) and laminin beta-2 (LAMB2) in Huh7 cells, which fits the previously observed interaction between DENV NS1 and LAMB1 in yeast two-hybrid assays^29^.

We also uncovered a network of enzymes participating in post-translational modifications in both cell types, suggesting that these proteins are *bona fide* interactors of NS1. With the exception of protein-lysine 6-oxidase (LOX) and peptidyl-prolyl cis-trans isomerase FKBP10 (FKBP10), all proteins in this network were prolyl 3-hydroxylases (P3H1, P3H2 and P3H3) or alpha subunits of prolyl 4-hydroxylase (P4HA1 and P4HA2). Prolyl 3-hydroxylases and prolyl 4-hydroxylases are mostly known to post-translationally modify collagens in the ER and are often referred to as collagen prolyl hydroxylases to distinguish them from the cytoplasmic P4H enzymes, which modify hypoxia-inducible factor (HIF-P4Hs)^30^. In the presence of oxygen, α-ketoglutarate, ferrous iron and ascorbate, collagen prolyl hydroxylases convert proline to (2S,3S)-hydroxyproline (prolyl 3-hydroxylases) or (2S,4R)-hydroxyproline (prolyl 4-hydroxylases)^20,31^, usually affecting stability of the modified protein^20,32^. Interestingly, the ER-resident chaperone FKBP10 (also known as FKBP65) is also required for collagen proteostasis^33-35^. While collagens are considered the main substrate for collagen prolyl 4-hydroxylases, other substrates have been reported for prolyl 4-hydroxylases, including the RNAi effector protein Argonaute 2 and the complement protein complex C1q ^21,36,37^.

The interaction between prolyl hydroxylases and NS1 prompted us to scrutinize our mass spectrometry data for direct evidence that NS1 contains hydroxyprolines. We focused on three residues, two containing the canonical XPA and XPG recognition sequences for collagen prolyl 4-hydroxylases^38^, residues P181 (tripeptide DPA) and P281 (tripeptide CPG), as well as the P267 residue embedded in a tripeptide that is highly conserved among diverse flaviviruses (Fig. 3a, Supplementary Table 7). In Supplementary Fig. 2, the MS/MS spectra for P181 and P267 containing fragments are shown. For residue P281, the fragmentation spectrum is shared with residue P267, has only a few annotated fragment ions and is therefore not shown. For residue P267 in NPCs, a fragmentation spectrum containing fragments that correspond to a hydroxyl group on P267 was observed in one of three replicates. In contrast, we did not observe a good fragmentation spectrum in Huh7 cells, and no modified proline reporter ions were identified in the MS/MS spectrum. For residue P181, a good fragmentation spectrum was obtained in Huh7 cells, which included a highly abundant Y-ion containing the modified proline residue. Together, these results suggest that ZIKV NS1 is a substrate for collagen prolyl 4-hydroxylases.

**Figure 3.**
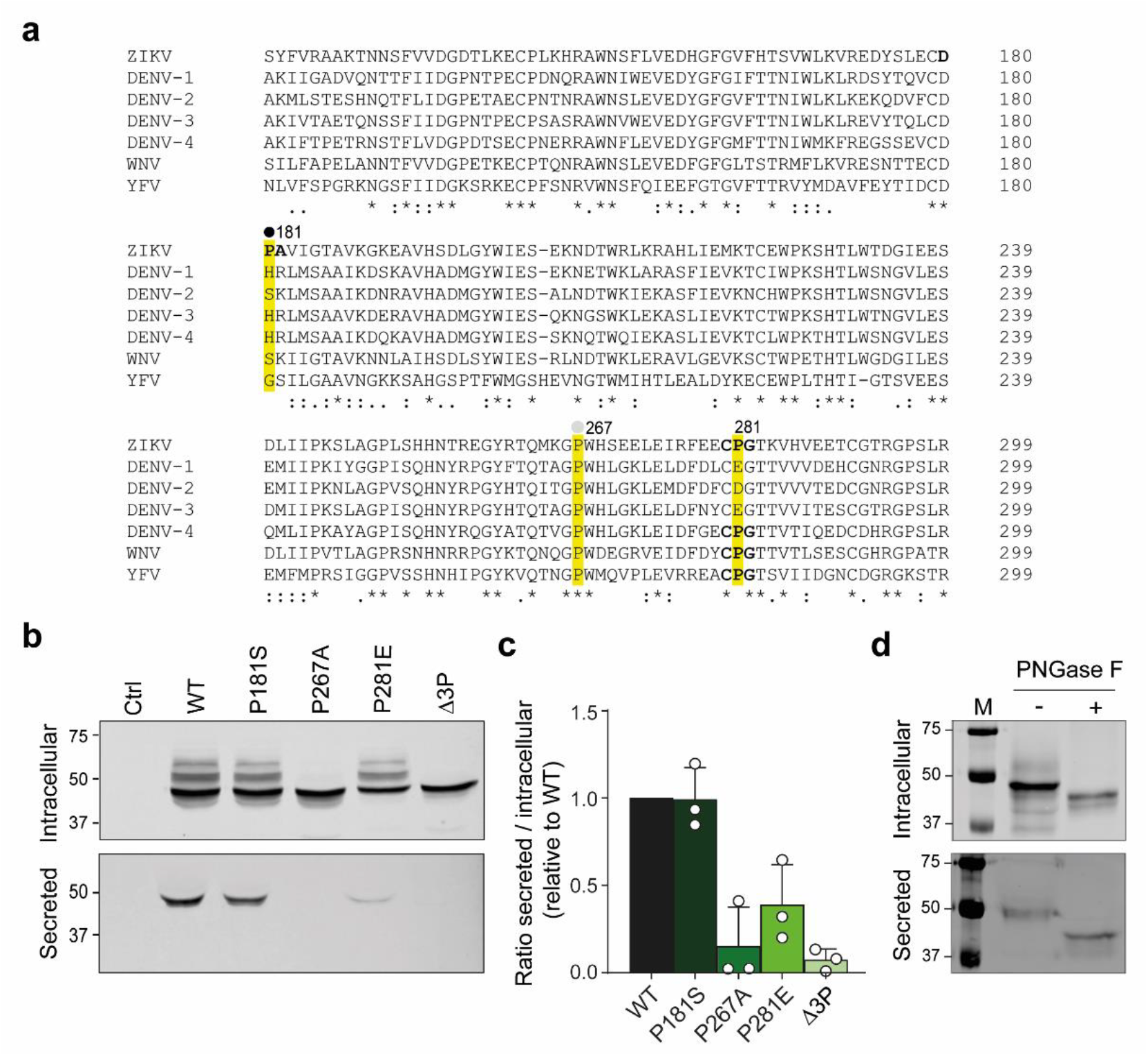
Proline residues are required for NS1 secretion. **a** Alignment of the NS1 sequence (amino acids 171-289) from ZIKV (NC_012532), dengue virus (DENV) serotypes 1–4 (NC_001477, NC_001474, NC_001475, NC_002640), West Nile virus (WNV, NC_009942) and yellow fever virus (YFV, NC_002031). Fully conserved residues are indicated by asterisks, conservation of strongly similar amino acids is indicated by colons, and conservation of weakly similar amino acids is indicated by dots. A hydroxyproline site in ZIKV NS1 with mass spectrometry support in multiple replicates is indicated with a closed circle and a site with support in one of three replicates is indicated with a grey circle. Canonical recognition sequences for collagen prolyl 4-hydroxylases are shown in bold. Proline residues selected for further investigation are highlighted in yellow. **b** Intracellular and secreted NS1 protein levels from HeLa cells transfected for 48 hours with the indicated wildtype (WT) or mutant NS1-FLAG expression constructs. Untransfected cells were taken along as a control (Ctrl). NS1 was detected using an antibody against FLAG. Numbers indicate marker sizes (kDa). Equal loading was confirmed with Ponceau staining (Supplementary Fig. 7a). **c** Quantification of the NS1 signal in the secreted over the intracellular fraction, normalized to WT NS1. Bars represent mean and SD of three independent experiments, open circles are the individual values. **d** Glycosylation status of ZIKV NS1. Hela cells were transfected with an expression plasmid encoding WT NS1-FLAG, and lysates from cells or culture supernatant were analyzed by western blot following treatment with (+) or without (-) PNGase F. The left lane is the size marker with sizes (kDa) indicated. A representative of two independent experiments is shown.

### NS1 proline mutants have maturation and secretion defects

We examined whether putative hydroxyprolines affect expression and trafficking of NS1 by mutating the corresponding proline residues in NS1 expression plasmids containing a C-terminal FLAG-tag. We informed our mutagenesis strategy using an NS1 alignment of ZIKV with other orthoflaviviruses, the four serotypes of dengue virus (DENV), West Nile virus (WNV) and yellow fever virus (Fig. 3a). P181 was replaced by serine, the corresponding residue in WNV and DENV-2, and P281 was replaced by glutamic acid, the corresponding residue in DENV-1 and DENV-3. P267 was replaced by alanine, as it was fully conserved in all flaviviruses analyzed, including insect-specific flaviviruses (Supplementary Fig. 3). In addition, we constructed an expression plasmid that contained all three mutations, referred to as the triple mutant or Δ3P.

We first analyzed intracellular NS1 expression and secretion of the mutants in HeLa cells, chosen for their higher transfection efficiency than Huh7 cells. Intracellular NS1 was detected as a dominant band and additional higher molecular weight bands on western blot, all of which were sensitive to PNGase F treatment, indicating that these represent N-glycosylated forms of NS1 (Fig. 3b-d), in line with previous observations^39,40^. For all mutants, intracellular NS1 expression was comparable to that of wildtype (WT) NS1, whereas NS1 secretion was almost completely abolished in the P267A and Δ3P mutants (Fig. 3b, c). Secretion of P218E NS1 was also strongly impaired, although less pronounced than observed for the P267A and Δ3P mutants, whereas the P181S mutant was not affected.

We next investigated whether the proline substitutions altered NS1 localization by confocal microscopy. For WT NS1 we observed a pattern consistent with a protein excreted through the secretory pathway with punctate intracellular staining, presumed secretory vesicles, and strong staining at the plasma membrane. The localization of P181S NS1 resembled the localization of WT NS1, although we observed some intracellular retention and slightly lower, albeit nonsignificant, membrane localization. All other mutants seemed to aggregate in tubular intracellular structures, presumably the ER, and were expressed at lower expression at the plasma membrane (Fig. 4a, b). Indeed, costaining with the calnexin ER marker and phalloidin to observe cell architecture confirmed that WT NS1 is mainly expressed at the plasma membrane, where it induces filipodia-like structures, whereas the P267A and D3P mutants were retained in the ER (Fig. 4c, d). Severity of the phenotype correlated with the maturation defects observed in western blot, with strong intracellular accumulation and low cell surface expression for P267A and Δ3P, and reduced NS1 expression at the plasma membrane for P281E.

**Figure 4.**
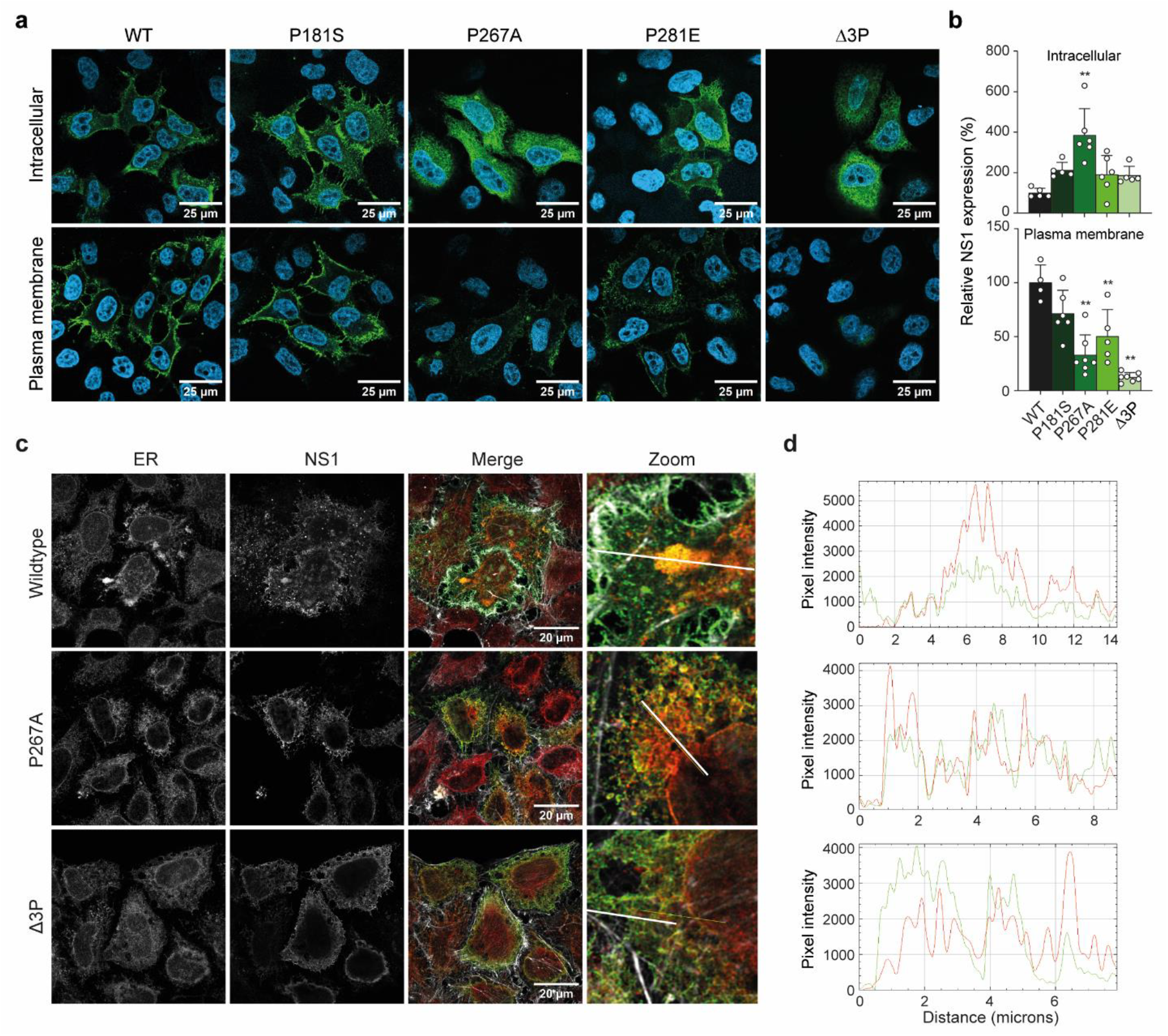
NS1 proline substitutions cause intracellular protein aggregation. **a** Confocal laser scanning microscopy images of HeLa cells transfected for 48 hours with WT or mutant NS1-FLAG expression constructs. Cells were fixed and either stained directly (plasma membrane staining) or stained after permeabilization (intracellular staining). NS1 was stained with antibodies against FLAG (green) and nuclei were visualized with Hoechst (blue). **b** Quantification of the NS1 signal in **a**, normalized to WT NS1 for each condition. Data were analyzed with one-way ANOVA with Dunnett’s post-hoc test (** *p* < 0.01). Bars represent the mean and SD of 24–28 positive cells for intracellular NS1 and 8 (triple mutant) or 24–31 positive cells (WT and other mutants) for plasma membrane localized NS1 per condition in a single experiment. Data from individual slides are shown as circles. **c** Confocal microscopy images of Hela cells expressing WT, P267 and D3P NS1-FLAG, stained with antibodies against FLAG and the calnexin ER marker. F-actin was stained with phalloidin. In the merged figure, NS1 is shown in green, ER in red, and F-actin in gray. **d** Intensity profiles of the NS1 (green) and ER (red) signal along the marked line in the zoomed image.

### P267A and Δ3P recombinant viruses have replication defects

To examine whether the NS1 proline substitutions affect ZIKV replication, we introduced these mutations into the viral genome using CPER cloning and analyzed viral titers. Ten days after transfection of Vero cells with the CPER reaction products (passage 0), viral titers of ZIKV^P181S^, ZIKV^P267A^ and ZIKV^P281E^ were more than one log lower than ZIKV^WT^ titers, whereas the ZIKV^Δ3P^ titers were undetectable (Fig. 5a). While these results suggest a replication defect for these mutants, this was not a controlled infection experiment, and the titers should not be interpreted as quantitative readout for replication efficiency. We used these passage 0 stocks to infect C6/36 cells and obtained roughly similar titers for the resulting passage 1 (p1) viruses for all mutants. However, Sanger sequencing of the region spanning the NS1 mutation in the p1 viruses revealed that ZIKV^P267A^ and ZIKV^Δ3P^ reverted back to WT sequences, indicating that these mutations are deleterious to viral replication, whereas the other two mutations were retained in the p1 stocks.

**Figure 5.**
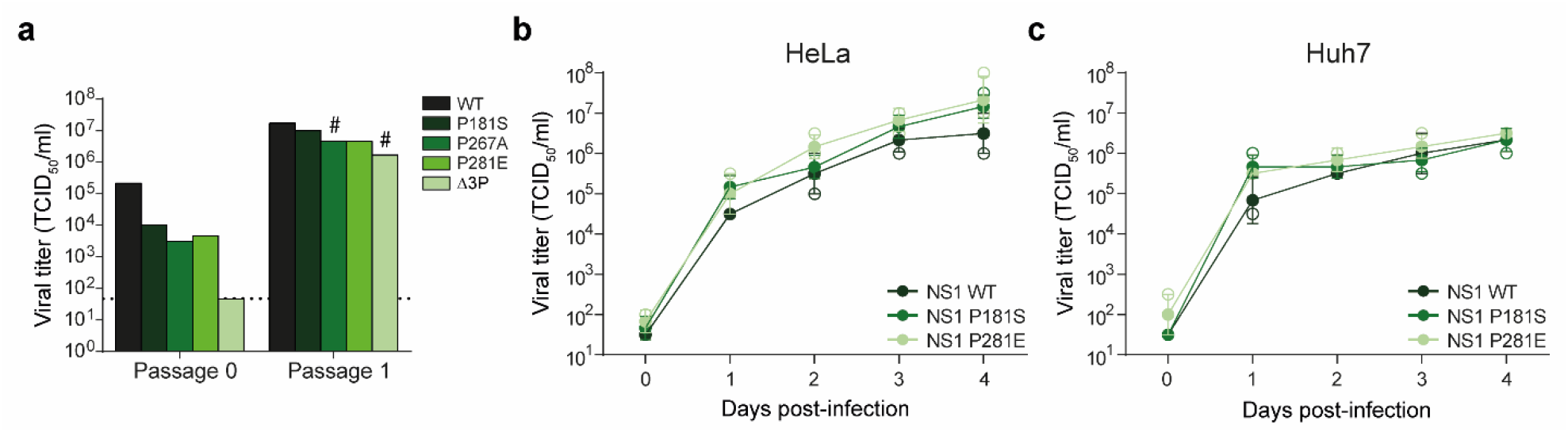
Replication kinetics of proline substituted ZIKV. **a** Infectious titers in the culture supernatant of Vero cells at 10 days after transfection with CPER products (passage 0) or C6/36 cells at 96 hours post-infection with ZIKV^WT^ or the single proline mutant viruses at an MOI of 0.001 (passage 1). For ZIKV^Δ3P^, passage 0 titers were undetectable, and a maximum volume of 1 ml was used to inoculate C6/36 cells. ZIKV^P267A^ and ZIKV^Δ3P^ viruses reverted back to WT sequences at passage 1 (indicated with a hash). **b, c** Infectious titers in the culture supernatant of HeLa (**b**) or Huh7 (**c**) cells at the indicated time points post-infection with ZIKV^WT^ or the indicated NS1 mutant viruses at an MOI of 0.1. Data were log-transformed and analyzed with one-way ANOVA with Dunnett’s post-hoc test, but differences between viruses were not statistically significant. Line graphs represent mean (filled symbols) and SD of two biological replicates. Open symbols represent individual data points.

We next analyzed the replication kinetics of ZIKV^P181S^, ZIKV^P281E^ and ZIKV^WT^ in HeLa and Huh7 cells over the course of four days. There were no significant differences in titers between the mutants and ZIKV^WT^ in either cell line, indicating that the P181S and P281E substitutions do not affect viral replication (Fig. 5b, c), despite the NS1 secretion defect observed especially for NS1 P281E (Fig. 3b). These results suggest that these mutations do not impair the role of NS1 in viral replication but likely affect other NS1 functions, particularly those associated with its secreted form.

### Prolyl hydroxylases are essential for NS1 intracellular trafficking

Due to its unique cyclic structure, proline plays an important role in protein structure. Proline substitutions could therefore have affected NS1 folding, which may explain the observed maturation and secretion phenotypes (Fig. 3, Fig. 4). To confirm our results, we used a pharmacological inhibitor of P4H enzymes, ethyl 3,4-dihydroxybenzoate (EDHB), to analyze the effect of prolyl hydroxylation on intracellular trafficking of WT NS1. EDHB is commonly used to inhibit collagen prolyl 4-hydroxylases, although it inhibits cytoplasmic HIF-P4H enzymes as well^41,42^.

Hela cells were transfected with the NS1-FLAG expression construct, treated with EDHB, and NS1 localization was analyzed using confocal microscopy. In the DMSO-treated cells, intracellular NS1 was especially abundant at or in close proximity to the plasma membrane and at cell-cell contact sites (Fig. 6a, b). In EDHB-treated cells, intracellular NS1 was retained throughout the cytoplasm, suggesting a defect in maturation and/or secretion. As a consequence, the intracellular NS1 signal intensity was increased in EDHB treated cells, which was especially pronounced at the higher concentrations (Fig. 6b). Moreover, at the highest concentration tested, EDHB treatment resulted in enlarged cells (Fig. 6a), which was reflected by an increased area of the intracellular NS1 signal (Supplementary Fig. 4). In conclusion, these data indicate that pharmacological inhibition of P4Hs impairs intracellular trafficking of NS1.

**Figure 6.**
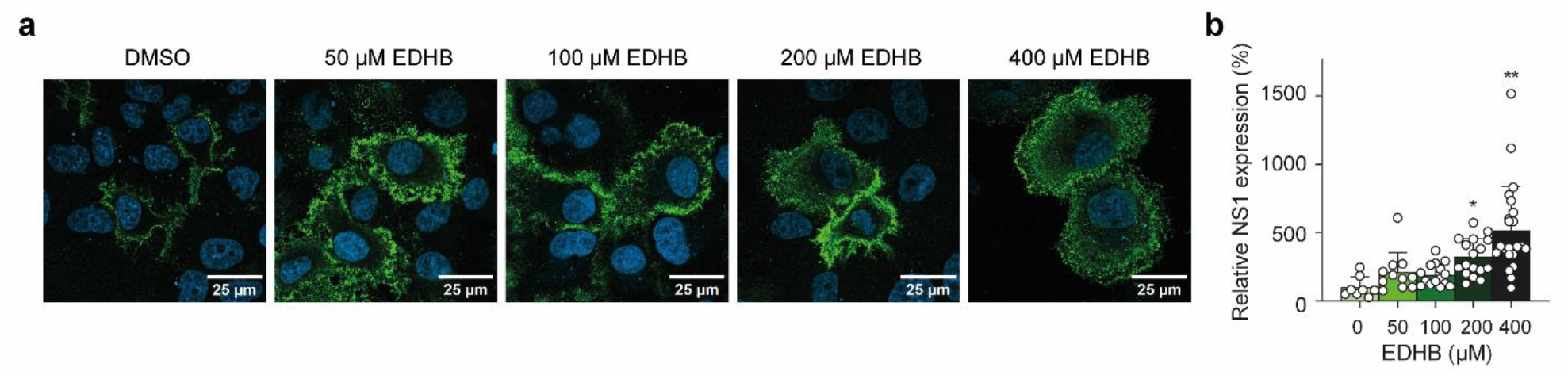
Prolyl hydroxylation is required for intracellular NS1 trafficking. **a** Confocal laser scanning microscopy images of HeLa cells expressing NS1-FLAG, treated with the indicated concentrations of EDHB or DMSO as a control. Cells were transfected for 48 hours and treated with EDHB or an equivalent volume of DMSO from 5 hours post-transfection onwards. Cells were fixed and permeabilized, and NS1 was stained with FLAG antibodies (green) and nuclei were visualized with Hoechst (blue). **b** Signal quantification of the NS1 staining of panel **a**, normalized to the DMSO control. Data were analyzed with one-way ANOVA with Dunnett’s post-hoc test (* *p* < 0.05; ** *p* < 0.01). Bars represent the mean and SD of 39–58 positive cells per condition. Individual data points are shown as circles.

### Inhibition of prolyl hydroxylases reduces ZIKV replication

Finally, we examined whether prolyl hydroxylation is important for viral replication by treating ZIKV^NS1:FLAG^ infected HeLa cells with EDHB. We first analyzed viability of uninfected EDHB-treated cells, to ensure that potential EDHB-induced cytotoxicity would not confound the infection experiments. EDHB concentrations of 400 µM or higher were cytotoxic, reducing relative cell viability to < 70%, whereas treatment with 100 or 200 µM EDHB reduced viability to just above 70% (Fig. 7a). In contrast, EDHB concentrations up to 50 µM did not affect cell viability and were consequently used in the infection experiments. Treatment with 25 or 50 µM EDHB decreased NS1 protein expression by 56–81% at 48 hours post-infection (Fig. 7b). Viral RNA levels were significantly reduced by 39–53% (*p* < 0.01) and viral titers were reduced more than half a log when cells were treated with 50 µM EDHB, although this difference did not reach statistical significance (Fig. 7c, d). Finally, we assessed whether EDHB also affected ZIKV replication at a later time point. The effects on NS1 expression, ZIKV RNA levels and viral titers were similar at 48 and 96 hours post-infection, although the differences in viral RNA levels and titers were only statistically significant for 50 µM EDHB treatment at 48 hours post-infection (Supplementary Fig. 5). Together, these results are in line with observations by Aviner et al. and indicate that P4H activity is required for efficient ZIKV replication^43^.

**Figure 7.**
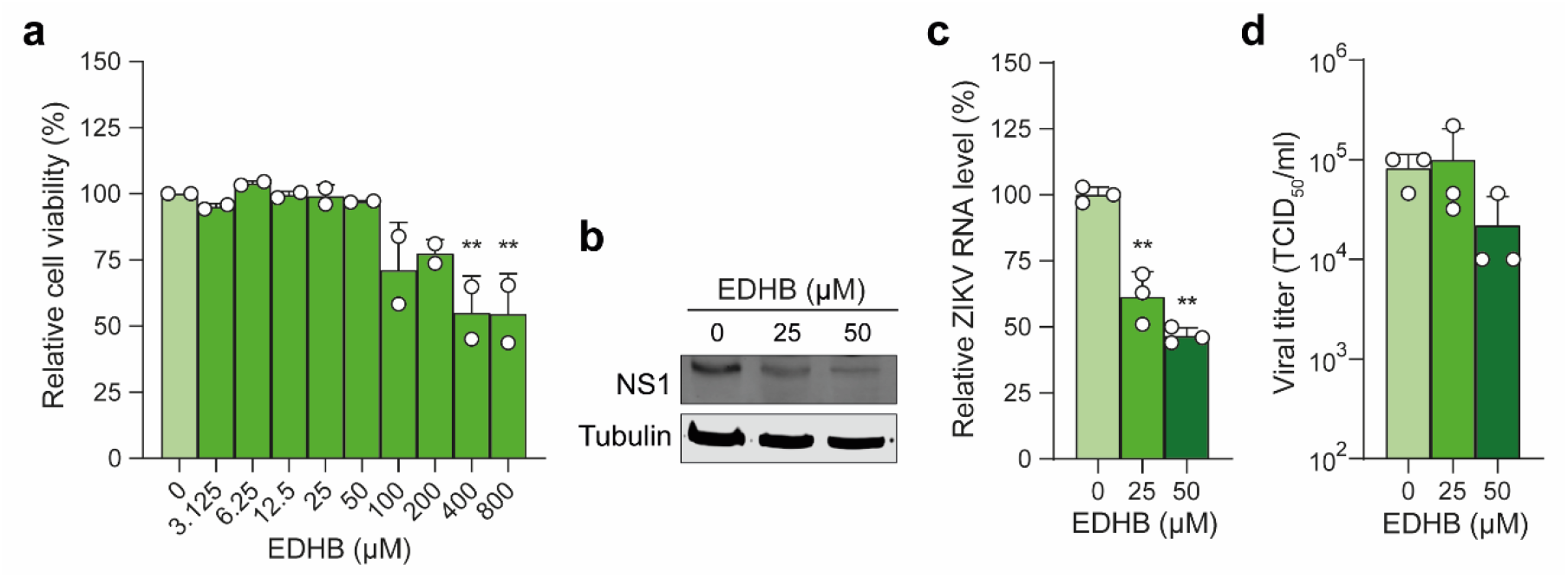
Prolyl hydroxylase inhibitors reduce ZIKV replication. **a** Cell viability assay of HeLa cells treated with the indicated concentrations of EDHB for 48 hours. The cell viability values were normalized to the DMSO treated control. **b–e** Intracellular NS1 protein levels (**b**), ZIKV RNA levels (**c**), and infectious titers in the culture supernatant of HeLa cells (**d**) at 48 hours post-infection. Cells were infected with ZIKV^NS1:FLAG^ at an MOI of 0.01 and treated with 25 or 50 µM EDHB or DMSO from 2 hours post-infection onwards. Intracellular NS1 protein was detected using a FLAG antibody. Tubulin was used as a loading control. For panel **a**, bars and dots represent mean +/ SEM of two independent experiments, each performed with three replicate wells. Dots represent the mean of each experiment. For panels **c** and **d**, bars and dots represent mean +/-SD and individual data of three replicate wells, with dots representing the individual data points. Data were analyzed with one-way ANOVA with Dunnett’s post-hoc test, using log-transformed data for panel **d** (** *p* < 0.01).

## Discussion

Viral proteins interact with host proteins to facilitate viral replication and antagonize antiviral responses. Identifying these host factors can therefore provide valuable insights into the molecular mechanisms underlying these processes. Using quantitative mass spectrometry on ZIKV-infected cells, we identified collagen P4H enzymes as interactors of ZIKV NS1 and found potential hydroxyproline residues in NS1. Moreover, results from pharmacological inhibition establish P4H enzymes as important ZIKV host factors affecting NS1 maturation and secretion.

Collagen P4Hs were also identified by Aviner *et al*. as post-translational modifiers of multiple ZIKV and DENV proteins^43^. Although this study did not detect hydroxyprolines in NS1, reduced NS1 levels were observed in ZIKV-infected cells in which *P4HA1* and *P4HA2* were knocked down. In addition, viral titers of ZIKV and DENV were approximately two logs lower in *P4HA1* and *P4HA2* knockdown cells compared to control cells, corroborating the importance of prolyl hydroxylation for efficient viral replication. We hypothesize that prolyl hydroxylation influences folding, maturation and secretion of ZIKV NS1 dimers and hexamers. Together with the Aviner *et al*. study, our results identify proline hydroxylation as an essential post-translational modification of flavivirus proteins.

Treatment of ZIKV-infected cells with 25 or 50 µM EDHB resulted in a relatively modest replication phenotype in our experiments. The concentrations used in this study were relatively low compared to those reported in the literature (typically 200–800 µM^42,44-46^), as we observed cytotoxic effects at concentrations exceeding 50 µM. Notably, higher EDHB concentrations resulted in more pronounced NS1 trafficking defects in our microscopy analyses, which are expected to lead to stronger replication phenotypes. Although EDHB also leads to increased HIF-1α expression^47^, it seems unlikely that this is the reason for the antiviral effect that we observed, since HIF-1α was reported to enhance, rather than decrease, DENV RNA replication^48^.

We found that the ZIKV^P181S^ and ZIKV^P281E^ viruses did not exhibit a replication defect in two human cell lines. The high fitness of these viruses was somewhat surprising given the substantial reduction in NS1 cell surface expression and secretion, particularly for NS1 P281E. However, it remains possible that these mutations reduce viral fitness or pathogenicity *in vivo*. For example, loss of DENV NS1 glycosylation reduced complement binding and abolished NS1-induced endothelial hyperpermeability, two functions associated with the secreted form of NS1^49,50^.

Our proteomic study identified several additional NS1 interactors that we did not follow up on in this study. For example, our dataset includes three enzymes that are involved in attachment and further modification of O-linked glycans. While we cannot exclude that these proteins were indirect hits due to their common localization in the ER or the Golgi apparatus, it may be interesting to investigate whether ZIKV NS1 is O-glycosylated and, if so, whether it has functional consequences. So far, hepatitis C virus is the only virus within the *Flaviviridae* family for which O-glycosylation has been demonstrated^51,52^. However, because the prediction and identification of O-glycosylation are technically challenging, the possibility that this modification occurs in ZIKV or other orthoflaviviruses cannot be excluded^51^.

Several prior studies have used MS-based proteomics to identify interactors of NS1 of ZIKV and the related orthoflavivirus DENV (serotype 2)^15-19,53,54^. None of these studies identified prolyl hydroxylases as significant NS1 interactors, suggesting that the use of infectious ZIKV may have been instrumental as most studies used epitope-tagged DENV or ZIKV NS1 transgenes or NS1-tagged DENV replicon (Supplementary Table 2). Moreover, manual inspection of the data suggests little overlap between interactors in our study and studies using NS1 transgenes (Supplementary Table 2). Whether this is due to differences in the context in which NS1 is expressed, the use of different cell types and viruses, or different criteria for assigning a hit significant remains unclear (Supplementary Table 2). Despite these limitations, we note that several of our ZIKV NS1 interactors were found as DENV2 NS1 interactors in the study of Hafirassou *et al*.^16^ (Exportin-2, CSE1L/EXP2; 7-dehydrocholesterol reductase, DHCR7; NADH dehydrogenase [ubiquinone] 1 alpha subcomplex subunit 5, NDUFA5; Endophilin-B1, SH3GLB1; Large neutral amino acids transporter small subunit 1, SLC7A5), which would be promising candidates as pan-flavivirus host factors.

Our study has several limitations. First, we retrieved FLAG:NS1 after cell lysis of infected cells without prior cross-linking. We can thus not formally exclude that protein-protein interactions have occurred after cell lysis and may be non-physiological. Additional NS1 interactors need confirmation and functional validation in follow-up studies. Second, while proline mutations altered NS1 trafficking and secretion and chemical inhibition of prolyl hydroxylases induced similar defects in WT NS1, we did have not directly demonstrated that the defects caused by the proline mutations are due to impaired prolyl hydroxylation. Third, while the main transmission route of ZIKV involves both a human host and a mosquito vector, our analyses focused exclusively on the role of prolyl hydroxylation in human cells. Given that prolyl hydroxylation is an evolutionary conserved posttranslational modification^55,56^ and that *Aedes* mosquitoes encode orthologs of prolyl 3-hydroxylase and prolyl 4-hydroxylase genes, future studies should investigate whether prolyl hydroxylation also contributes to ZIKV replication and transmission in the mosquito vector.

In conclusion, our results indicate that prolyl hydroxylation is required for ZIKV NS1 maturation and secretion and that this posttranslational modification is required for efficient replication. It will be interesting to investigate whether proline hydroxylation is essential for viral replication of other viruses as well and whether it could be a target for the development of broadly acting antiviral drugs.

## Material and methods

### Cells and viruses

Vero FM (ATCC CCL-81), Vero E6 (ATCC CRL-1586), HeLa-R19 (RRID:CVCL_M763) and Huh7 (RRID:CVCL_0336) cell lines were cultured in Dulbecco’s modified Eagle medium (DMEM) with 4.5 g/L D-glucose (Life Technologies), supplemented with 10% heat-inactivated fetal calf serum (FCS; Sigma-Aldrich) and 50 U/ml penicillin and 50 μg/ml streptomycin (1% pen/strep; Life Technologies) at 37 °C and 5% CO_2_. C6/36 cells (ATCC CRL-1660) were maintained at 28 °C in Leibovitz’s L-15 medium (Life Technologies), supplemented with 10% FCS, 1× MEM non-essential amino acids (Life Technologies), 2% tryptose phosphate broth (Sigma-Aldrich) and 1% pen/strep. Healthy human donor-derived iPSCs containing a doxycycline-inducible rtTA/Ngn2 transgene^57^ were grown in Essential 8 Flex medium supplemented with 0.5 µg/ml puromycin (Sigma-Aldrich),50 µg/ml G418 (Sigma-Aldrich) and 1% pen/strep in 6-well plates coated with 0.5 µg/cm^2^ vitronectin (Life Technologies). Cells were passaged as clumps every 4–6 days using 0.5 mM EDTA in PBS^58^. iPSCs were differentiated into NPCs according to the monolayer protocol of the STEMdiff Neural System (STEMCELL Technologies). Briefly, iPSCs were seeded as single cells in 6-well plates coated with Matrigel (Corning Life Science) in complete Neural Induction Medium, consisting of STEMdiff Neural Induction Medium with SMADi Neural Induction Supplement (STEMCELL Technologies) and 1% pen/strep, which was supplemented with 1× RevitaCell (Life Technologies) only on the day of seeding. NPCs were passaged using Accutase (STEMCELL Technologies) when they reached 80-90% confluence. From passage 3 onwards, NPCs were cultured in STEMdiff Neural Progenitor Medium (STEMCELL Technologies) with 1% pen/strep (NPC medium). Experiments were performed with cells from passage four and five.

ZIKV strain H/PF/2013 was obtained from the European Virus Archive (EVAg). Unless otherwise stated, stocks were generated by infecting C6/36 cells at an MOI of 0.1 in complete Leibovitz’s L-15 medium. Upon observation of cytopathic effect (CPE), the cell culture supernatants were cleared by centrifugation for 5 minutes at 500 x *g*, aliquoted and stored at -80 °C. Viral titers were determined by end-point dilution assay as described previously^59^. The 50% tissue culture infective dose (TCID_50_) per ml was calculated according to the Reed and Muench method^60^.

### Expression plasmids

A ZIKV NS1 expression vector was generated by cloning the sequence spanning the last transmembrane domain of the E protein and the complete NS1 sequence as well as a C-terminal 3x FLAG-tag into the pEF-2K-NS4B-FLAG plasmid, encoding the DENV NS4B protein (kindly provided by Andrew Davidson, University of Bristol). First, an intermediate cloning plasmid was generated by replacing the existing 1× FLAG-tag and most of the sequence encoding the DENV proteins by a 3× FLAG tag by digesting the pEF-2K-NS4B-FLAG plasmid with HpaI (New England Biolabs, NEB) and EcoRI-HF (NEB). Oligonucleotides encoding a 3× FLAG-tag (Supplementary Table 1) were annealed and inserted in the digested plasmid as described previously^61^. The sequence encoding the last transmembrane domain of the ZIKV E protein and the full ZIKV NS1 protein was amplified from the pCCI-CMV-ZIKV WT IC plasmid^23^ using Phusion High-Fidelity DNA Polymerase (NEB). This insert was cloned into the intermediate plasmid using In-Fusion cloning (Takara Bio) after digesting the plasmid with HpaI and NcoI-HF (NEB) to generate pEF-ZIKV-NS1-FLAG.

The ZIKV NS1 mutant expression plasmids were made by site-directed mutagenesis on pEF-ZIKV-NS1-FLAG using the In-Fusion cloning kit with the primers in Supplementary Table 1. Two additional rounds of site-directed mutagenesis were performed to produce the triple proline mutant. All generated plasmids were validated with Sanger sequencing.

### Compounds

EDHB (CAS No. 3943-89-3) was purchased from Sigma-Aldrich. The compound was dissolved in DMSO to a final concentration of 1000 mM and stored at -20 °C as single use aliquots.

### Circular polymerase extension reaction cloning

ZIKV^NS1:FLAG^, ZIKV^WT^, and proline mutant viruses were produced with a circular polymerase extension reaction (CPER) according to the protocol of Slonchak et al.^62^ using the pCCI-SP6-ZIKV wild-type infectious clone (ZIKV WT IC, ^23^) and mutant NS1 expression plasmids as templates. A pUCIDT plasmid (Integrated DNA Technologies) containing the UTR-linker sequence, which consists of the last 22 nucleotides of the ZIKV WT IC 3’ UTR, a hepatitis delta virus ribozyme, a CMV promoter, and the first 22 nucleotides of the ZIKV WT IC 5’UTR was purchased from Integrated DNA Technologies (Supplementary Fig. 6). The CPER fragments were generated by PCR with Phusion polymerase (New England Biolabs) using the linker-containing plasmid and the ZIKV WT IC or the mutant NS1 expression plasmids as templates with the primers listed in Supplementary Table 1. The FLAG-tag was inserted into the CPER product by placing the FLAG-tag sequence in the forward primer for the second fragment (2989 bp) and the reverse primer of the first fragment (3537 bp) (see also Fig. 1a). Equimolar amounts of each PCR fragment were used in a CPER reaction using the Primestar GXL polymerase (Takara Bio) as described ^62^.

Fifty µl of the CPER products were transfected into Vero FM cells using Lipofectamine LTX with Plus Reagent (Invitrogen). The medium was replaced with DMEM supplemented with 2% FCS and 1% pen/strep the next day and passage 0 stocks were collected at 10 days post-transfection. Passage 1 and 2 stocks were generated by inoculating C6/36 cells at an MOI of 0.1 for the viruses used for mass spectrometry or an MOI of 0.001 for the single proline mutant viruses and the wildtype virus that were grown in parallel. For the Δ3P mutant, 1 ml of passage 0 stock was used to inoculate C6/36 cells. Cell culture supernatants were collected upon appearance of CPE. The presence of the FLAG-tag and introduced mutations were verified by Sanger sequencing after the first and second passage for ZIKV^NS1:FLAG^ or after transfection and the first passage for the proline mutants. Passage 1 stocks were used for all experiments except mass spectrometry, for which passage 2 stocks were used.

### RNA isolation, reverse transcription and RT-qPCR

Total RNA was isolated with RNA-Solv reagent (Omega Biotek) according to the manufacturer’s instructions. RNA was treated with DNase I (Roche) before cDNA synthesis using TaqMan Reverse Transcription Reagents with random hexamers (Invitrogen) according to the manufacturers’ instructions. Viral RNA levels and expression of host genes were measured with RT-qPCR on a LightCycler 480 (Roche) using the GoTaq qPCR master mix (Promega) and the primers listed in Supplementary Table 1. Data were normalized to β-actin values and relative expression was calculated using the 2^-ΔΔCT^ method^63^.

### Immunoprecipitation of ZIKV NS1-FLAG for interactome analysis

Huh7 cells were seeded in eight T175 flasks at a density of 1.33 × 10^7^ cells/flask in complete DMEM and NPCs were seeded in six Matrigel-coated T175 flasks at a density of 4.0 × 10^6^ cells/flask in complete NPC medium. The cells were infected the following day with ZIKV^WT^ or ZIKV^NS1:FLAG^ at an MOI of 1.0 in DMEM without supplements for Huh7 cells or complete NPC medium for NPCs. The medium was refreshed 2 hours post-infection with DMEM containing 2% FCS and 1% pen/strep or complete NPC medium for Huh7 cells and NPCs, respectively. For the NPCs, half of the medium was replaced with fresh NPC medium one day post-infection. At 48 hours post-infection, the cell culture supernatant was stored at -80°C and the cells were detached with trypsin (Huh7 cells) or Accutase (NPCs). Complete DMEM (Huh7 cells) or DMEM/F-12 (NPCs) was added to the cell suspensions, which were subsequently centrifuged for 5 minutes at 900 × *g* at 4°C. After centrifugation, the pellets were washed twice in PBS by centrifugation for 2 minutes at 500 × *g* at 4°C. The cells were lysed in lysis buffer (10 mM Tris/HCl pH 7.4, 150 mM NaCl, 0.5 mM EDTA, 0.5% NP-40 and 1× cOmplete protease inhibitor [Roche]) for one hour at 4°C with end-over-end rotation. Lysates were cleared by centrifugation at 16,000 × *g* for 30 minutes at 4 °C and the protein concentration in the lysates was determined using a BCA assay (Pierce) according to the manufacturer’s instructions.

Protein concentrations of ZIKV^WT^ or ZIKV^NS1:FLAG^ infected cell lysates were equalized using dilution buffer (10 mM Tris/HCl pH 7.4, 150 mM NaCl, 0.5 mM EDTA and 1× cOmplete protease inhibitor) and 0.825 mg (Huh7 cells) or 1.0 mg (NPCs) of each lysate was mixed with 10 µl anti-FLAG M2 affinity gel (Sigma-Aldrich), which was equilibrated in dilution buffer before use. The samples were incubated overnight at 4°C with end-over-end rotation. The beads were collected by centrifugation for 1 minute at 1016 x *g* at 4°C, washed thrice with dilution buffer and once with PBS. The beads were resuspended in PBS, transferred to fresh tubes, and centrifuged for 1 minute at 1016 x *g* at 4°C. The PBS was removed, and the dry beads were stored at -20°C until further processing for mass spectrometry.

### Mass spectrometry and protein-protein interaction analysis

Proteins were on-bead digested as follows: 50 µl elution buffer (2M urea, 100 mM Tris-HCl pH 8.0, 10 mM DTT) was added and incubated at 25 °C while shaking. Next, iodoacetamide was added to a final concentration of 50 mM and incubated for 10 minutes in the dark. Then, 0.25 µg of MS-grade trypsin (Promega) was added and incubated at 25 °C for 2 hours while shaking. The supernatants were collected, and the remaining peptides were collected from the beads by an addition wash with 50 µl elution buffer. The first supernatant of each sample was combined with the wash supernatant and further incubated overnight at 25 °C in the presence of 0.1 µg fresh trypsin. The next day, samples were acidified using 10 µl 10% TFA and loaded on stagetips (Rappsilber) and stored until MS analysis.

Peptides were analyzed during a 60-minute reverse phase gradient on a Thermo Easy-nLC1000 with a 30 cm C18 ESI column connected online to a Thermo LTQ Exploris 480 operated in Top20 mode with a dynamic exclusion of 45 seconds. RAW data were analyzed using MaxQuant version 1.6.6.0^64^ and a human database (downloaded in June 2017) supplemented with the sequences of ZIKV proteins. Deamidation was added as a variable modification, and Label Free Quantification (LFQ) and iBAQ were enabled. For the analysis of hydroxyproline sites, MaxQuant was used with the same settings, only hydroxyproline was added as additional variable modification. Evidence for hydroxyproline was verified using the MaxQuant viewer. MaxQuant output was further processed using Perseus^65^. LFQ values were log_2_ transformed, and potential contaminants, reverse hits, and proteins with less than two peptides or without unique peptides were filtered out. Samples were grouped and only proteins with three valid values in at least one group were retained (assuming that specifically enriched proteins may not be identified in the negative control samples). Missing values were imputed, and a 2-samples *t*-test was performed. Volcano plots were generated using R. FC cut-offs were chosen in such a way that a minimal number of outliers was present in the negative control samples (ZIKV^WT^), whereas a maximum number of true interactors were enriched in the ZIKV^NS1:FLAG^ samples. Depending on the background, these cut-offs may vary between experiments.

Protein-protein interaction networks were analyzed using the STRING database^66^. Only protein-protein interactions with an interaction score of > 0.4 that were predicted to interact based on homology, co-expression, experimentally determined interaction, database annotation, and text mining as interaction sources were considered. Proteins without any predicted interactions were excluded and networks were visualized in Cytoscape^67^.

### Cell viability assay

HeLa-R19 cells were seeded in white 96-well plates at a density of 1 × 10^4^ cells/well. One day after seeding, EDHB or an equivalent volume of DMSO was added to the cells, and cell viability was determined 48 hours post-treatment with the CellTiter-Glo 2.0 Cell Viability Assay (Promega) using the Victor3 microplate reader (Perkin Elmer).

### Growth curve proline mutants

Hela-R19 and Huh7 cells were seeded in 6-well plates at a density of 5 × 10^5^ cells/well. Six hours after seeding, the cells were infected with ZIKV^WT^ or the proline mutants at an MOI of 0.1 in DMEM without supplements. At 2 hours after inoculation, the cells were washed with PBS and complete DMEM was added. Cell culture supernatants were collected at 0, 24, 48, 72 and 96 hours post-infection and titrated on Vero E6 cells.

### Western blot

For western blot analysis of transfected cells, HeLa-R19 cells were seeded in two wells of a 6-well plate per condition at a density of 375,000 cells/well in complete DMEM. The next day, the cells were mock-transfected or transfected with an NS1 expression plasmid using Fugene 6 (Promega) according to the manufacturer’s instructions. Approximately 48 hours post-transfection, the culture supernatant of each condition was collected, centrifuged for 5 minutes at 140 x *g*, and mixed with an equal volume of cold 2× Triton buffer (PBS with 0.5% Triton, 20% glycerol and 2× cOmplete protease inhibitor). Cells were washed with PBS, trypsinized and resuspended in complete DMEM. Cells were centrifuged for 5 minutes at 140 x *g* and the pellets were washed twice in cold PBS by centrifugation for 2 minutes at 140 x *g*. Cells were lysed in cold Triton buffer (PBS with 0.25% Triton X-100, 10% glycerol and 1× cOmplete protease inhibitor) for one hour at 4°C with end-over-end rotation. Lysates were cleared by centrifugation at > 16,000 × *g* for 30 minutes at 4°C and protein concentrations in the lysate and culture supernatants were determined using a BCA assay (Pierce) according to the manufacturer’s instructions. Protein concentrations were equalized using Triton buffer, mixed with 2× SDS sample buffer (120 mM Tris-HCl pH 6.8, 4% SDS, 10% β-mercaptoethanol, 20% glycerol, 0.04% bromophenol blue) and the samples were heated for 10 minutes at 95°C. Samples were resolved on 12.5% polyacrylamide gels, equal loading was confirmed using a Ponceau S stain (Sigma-Aldrich), and resolved samples were wet-blotted to nitrocellulose membranes. Membranes were blocked for 30 minutes at room temperature in PBS with 5% milk and subsequently incubated overnight at 4°C with mouse anti-FLAG M2 antibody (Sigma-Aldrich, 1:1000 in PBS with 2.5% milk and 0.1% Tween-20). The following day, the membranes were washed three times with PBS with 0.1% Tween-20 (0.1% PBS-T) and incubated with IRDye 680 goat anti-mouse antibody (Li-Cor, 1:10,000 in PBS with 2.5% milk, 0.1% Tween-20 and 0.01% SDS) for 45 minutes at room temperature in the dark. Next, membranes were briefly washed thrice in PBS, followed by two washes with 0.1% PBS-T for 10 minutes and again briefly washed thrice in PBS. Membranes were imaged using the Odyssey-CLx Imaging System (Li-Cor), and signal intensity was quantified in Fiji as the raw integrated density in regions of interest after subtraction of the signal in a background region^68^. N-glycosylation was assessed by treatment of cell lysates of WT NS1-FLAG transfected cells with PNGase F (New England Biolabs) according to the manufacturer’s instructions under denaturing reaction conditions, followed by western blotting as described above.

For western blot analysis of infected cells, HeLa-R19 cells were seeded in a 6-well plate at a density of 5 × 10^5^ cells/well. The next day, the cells were infected with ZIKV^NS1:FLAG^ at an MOI of 0.01 in DMEM without supplements. After 2 hours, the medium was replaced by supplemented DMEM containing 25 or 50 µM EDHB, or an equivalent volume of DMSO. At 48 and 96 hours post-infection, the culture supernatant was collected for titration. In addition, ZIKV^NS1:FLAG^ infected cells were lysed in 1x Laemmli Sample Buffer (Bio-Rad). Samples were denatured for 10 minutes at 95°C and resolved on a 12.5% SDS PAGE gel, followed by semi-dry blotting on a 0.2 µm nitrocellulose membrane (Bio-Rad). Membranes were blocked, incubated with mouse anti-FLAG M2 (1:1000, Sigma) or rat anti-Tubulin (1:1000, Bio-Rad), washed, incubated with secondary antibodies goat anti-mouse IRDye-680RD or goat anti-rat IRdye-800CW (both 1:15000, Li-Cor) and washed again. Membranes were imaged using the Odyssey-CLx Imaging System and signal intensity was quantified using ImageJ. Uncropped western blot membranes and Ponceau stains are shown in Supplementary Figure 7.

### Confocal microscopy

HeLa-R19 cells were seeded on coverslips in 24-well plates at a density of 3.75 × 10^4^ cells/well in complete DMEM. The cells were transfected with 500 ng of wildtype or mutant NS1 expression plasmid using Fugene 6 according to the manufacturer’s instructions. At 5 hours post-transfection, the medium was replaced with complete DMEM or with complete DMEM supplemented with EDHB at the indicated concentrations or equivalent volumes of DMSO. Approximately 48 hours post-transfection, the cells were washed once with PBS and fixed in 4% PFA for 15 minutes.

Fixed cells were processed for immunofluorescence microscopy using two different protocols. For intracellular staining, cells were permeabilized in PBS with 0.5% Triton X-100 (v/v) for 20 minutes, washed twice with PBS with 0.05% Tween-20 (v/v, 0.05% PBS-T), and subsequently incubated in detergent-containing blocking buffer (PBS with 2% bovine serum albumin [w/v, BSA, Sigma-Aldrich], 2% normal goat serum [NGS], 0.1% Triton X-100, and 0.05% Tween-20) with 100 mM glycine for 30 minutes. For plasma membrane staining, the cells were not permeabilized and instead blocked with detergent-free blocking buffer (PBS with 2% BSA and 2% NGS) and 100 mM glycine for 30 minutes.

Mouse anti-FLAG M2 (Sigma-Aldrich, 1:1000) and Alexa Fluor 488 goat anti-mouse antibody (Invitrogen, 1:400) were diluted in detergent-containing or detergent-free blocking buffer and incubated on the coverslips for one hour. Coverslips were washed thrice with PBS for plasma membrane staining and with 0.05% PBS-T for intracellular staining between antibody incubations. Following secondary antibody incubation, the coverslips were washed twice in PBS for the plasma membrane staining or twice in 0.05% PBS-T followed by two PBS washes for the intracellular staining. Nuclei were stained using 5 µg/ml Hoechst, washed once with PBS, washed once with 0.5× PBS and thrice with water before mounting the coverslips on slides with Mowiol (Omnilabo). Images were acquired using a 63× oil immersion objective and a 40× water immersion objective on Zeiss LSM900 laser scanning microscope with Airyscan, and images were analyzed using ImageJ. A Li (NS1 mutants) or Otsu (EDHB treatment) threshold was applied to the images to create a cell mask and signal intensity within the mask was calculated using the integrated density value. For the EDHB treatment experiment, the integrated density value was divided by the number of cells to account for differences in cell numbers between images. After the measurements, the images were scaled using bilinear interpolation.

To analyze NS1 colocalization with the calnexin ER marker, HeLa-R19 cells were seeded on coverslips in 24-well plates in complete DMEM. The cells were transfected with 500 ng of wildtype, P267A or D3P mutant NS1 expression plasmid using XtremeGENE (Sigma-Aldrich) according to the manufacturer’s instructions. Approximately 24 hours post-transfection, the cells were washed once with PBS and fixed in 4% PFA for 15 minutes. Fixed cells were permeabilized with PBS with 0.1% Triton X-100 (v/v; PBS-T) for 10 minutes, washed once with PBS-T, followed by incubation with 0.1 M glycine for 7 minutes. Cells were washed once with PBS-T and subsequently blocked with PBS-T containing 2% NGS for 30 minutes. Mouse anti-Flag M2 (Sigma-Aldrich, 1:1000) and rabbit anti-calnexin antibodies (Proteintech #10427-2-AP, 1:400) were diluted in PBS-T containing 2% NGS and coverslips were incubated for 1 hour at room temperature. The coverslips were subsequently washed three times for 5 minutes in PBS-T and incubated with secondary antibodies goat anti-mouse Alexa Fluor 488 and goat anti-rabbit Alexa Fluor 594 diluted in PBS-T with 2% NGS (1:500) for 45 minutes, followed by three washes with PBS-T. Nuclei and F-actin were stained with DAPI (1:000 in PBS) and Phalloidin (Biotium, 1:200 in PBS) for 20 minutes. Coverslips were washed three times for 5 minutes with PBS and once with water before mounting on slides with ProLong Glass Antifade Mountant (Invitrogen). Images were acquired using a 63x oil immersion objective on a Zeiss LSM900 laser scanning microscope with Airyscan, and images were analyzed using ImageJ.

### Statistical analyses

Data were analyzed using GraphPad Prism 6.01 software. Viral titration data were log-transformed prior to statistical testing. Statistical significance of comparisons between two independent groups was tested with Student’s *t*-tests. For all other comparisons, one-way ANOVA was used, with correction for multiple testing. When one control group was compared to multiple treatments, Dunnett’s post-hoc test was used. In experiments with more than one control group, the Bonferroni correction was applied. When every mean was compared with every other mean in the same time point, the Holm-Šidák method was used.

## Data availability

Proteomics data have been deposited to the ProteomeXchange Consortium via the PRIDE partner repository with the dataset identifier PXD041488 ^69^. Other data generated and analyzed in this study are included in the published article and its supplementary information files.

## Acknowledgments

We thank Benoit Besson, Ezgi Taşköprü and other former and current members of the laboratory for discussions. We thank the Radboudumc Microscopic Imaging Center for support with fluorescence microscopy. We thank Andres Merits (University of Tartu, Tartu, Estonia) for kindly providing the Zika virus infectious clone, Nael Nadif Kasri (Radboudumc, Nijmegen, the Netherlands) for providing iPSCs, and Yin Xiang Setoh and Alexander Khromykh (University of Queensland, Brisbane, Australia) for advice on the CPER approach. ZIKV isolate H/PF/2013 was kindly provided by European Virus Archive (EVAg). This work was financially supported by a PhD grant from Radboud University Medical Center to RPvR and a PhD stipend from Radboud University Medical Center to MS. The Vermeulen laboratory is part of the Oncode Institute, which is partly funded by the Dutch Cancer Society. The funder played no role in study design, data collection, analysis and interpretation of data, or the writing of this manuscript.

## Author contributions

BB conceptualized the project, performed experiments, analyzed the data, prepared figures, and wrote the manuscript. GJO performed experiments and analyzed the data. MS performed microscopy analyses. CGS performed mass spectrometry and analyzed and interpreted the data. MV analyzed and interpreted the mass spectrometry data. PM provided supervision. RPvR conceptualized the project, acquired funding, interpreted the data, supervised the project, and wrote the manuscript. All authors reviewed the manuscript.

## References

1 Wikan, N. & Smith, D. R. Zika virus: History of a newly emerging arbovirus. The Lancet Infectious Diseases 16, e119–e126; 10.1016/S1473-3099(16)30010-X (2016).

2 Barrows, N. J. et al. Biochemistry and Molecular Biology of Flaviviruses. Chemical Reviews 118, 4448–4482; 10.1021/acs.chemrev.7b00719 (2018).

3 Rastogi, M., Sharma, N. & Singh, S. K. Flavivirus NS1: a multifaceted enigmatic viral protein. Virology Journal 13, 131; 10.1186/s12985-016-0590-7 (2016).

4 Muller, D. A. & Young, P. R. The flavivirus NS1 protein: Molecular and structural biology, immunology, role in pathogenesis and application as a diagnostic biomarker. Antiviral Research 98, 192–208; 10.1016/j.antiviral.2013.03.008 (2013).

5 Smith, G. W. & Wright, P. J. Synthesis of Proteins and Glycoproteins in Dengue Type 2 Virus-infected Vero and Aedes albopictus Cells. Journal of General Virology 66, 559–571; 10.1099/0022-1317-66-3-559 (1985).

6 Winkler, G., Maxwell, S. E., Ruemmler, C. & Stollar, V. Newly Synthesized Dengue-2 Virus Nonstructural Protein NS1 Is a Soluble Protein but Becomes Partially Hydrophobic and Membrane-Associated after Dimerization. Virology 171, 302–305; 10.1016/0042-6822(89)90544-8 (1989).

7 Winkler, G., Randolph, V. B., Cleaves, G. R., Ryan, T. E. & Stollar, V. Evidence that the Mature Form of the Flavivirus Nonstructural Protein NS1 is a Dimer. Virology 162, 187–196; 10.1016/0042-6822(88)90408-4 (1988).

8 Akey, D. L. et al. Flavivirus NS1 crystal structures reveal a surface for membrane association and regions of interaction with the immune system. Science 343, 881–885; 10.1126/science.1247749 (2014).

9 Watterson, D., Modhiran, N. & Young, P. R. The many faces of the flavivirus NS1 protein offer a multitude of options for inhibitor design. Antiviral Research 130, 7–18; 10.1016/j.antiviral.2016.02.014 (2016).

10 Noisakran, S. et al. Association of dengue virus NS1 protein with lipid rafts. Journal of General Virology 89, 2492–2500; 10.1099/vir.0.83620-0 (2008).

11 Westaway, E. G. & Goodman, M. R. Variation in Distribution of the Three Flavivirus-specified Glycoproteins Detected by Immunofluorescence in Infected Vero Cells. Archives of Virology 94, 215–228; 10.1007/BF01310715 (1987).

12 Gutsche, I. et al. Secreted dengue virus nonstructural protein NS1 is an atypical barrel-shaped high-density lipoprotein. Proceedings of the National Academy of Sciences 108, 8003–8008; 10.1073/pnas.1017338108 (2011).

13 Avirutnan, P. et al. Antagonism of the complement component C4 by flavivirus nonstructural protein NS1. Journal of Experimental Medicine 207, 793–806; 10.1084/jem.20092545 (2010).

14 Puerta-Guardo, H. et al. Flavivirus NS1 Triggers Tissue-Specific Vascular Endothelial Dysfunction Reflecting Disease Tropism. Cell Reports 26, 1598–1613.e1598; 10.1016/j.celrep.2019.01.036 (2019).

15 Coyaud, E. et al. Global interactomics uncovers extensive organellar targeting by Zika Virus. Molecular and Cellular Proteomics 17, 2242–2255; 10.1074/mcp.TIR118.000800 (2018).

16 Hafirassou, M. L. et al. A Global Interactome Map of the Dengue Virus NS1 Identifies Virus Restriction and Dependency Host Factors. Cell Reports 21, 3900–3913; 10.1016/j.celrep.2017.11.094 (2017).

17 Scaturro, P. et al. An orthogonal proteomic survey uncovers novel Zika virus host factors. Nature 561, 253–257; 10.1038/s41586-018-0484-5 (2018).

18 Shah, P. S. et al. Comparative Flavivirus-Host Protein Interaction Mapping Reveals Mechanisms of Dengue and Zika Virus Pathogenesis. Cell 175, 1931–1945; 10.1016/j.cell.2018.11.028 (2018).

19 Wang, Y., Uraki, R., Hwang, J. & Fikrig, E. TRiC/CCT Complex, a Binding Partner of NS1 Protein, Supports the Replication of Zika Virus in Both Mammalians and Mosquitoes. Viruses 12, 519; 10.3390/v12050519 (2020).

20 Gorres, K. L. & Raines, R. T. Prolyl 4-hydroxylase. Critical Reviews in Biochemistry and Molecular Biology 45, 106–124; 10.3109/10409231003627991 (2010).

21 Qi, H. H. et al. Prolyl 4-hydroxylation regulates Argonaute 2 stability. Nature 455, 421–424; 10.1038/nature07186 (2008).

22 Fulton, B. O., Sachs, D., Schwarz, M. C., Palese, P. & Evans, M. J. Transposon Mutagenesis of the Zika Virus Genome Highlights Regions Essential for RNA Replication and Restricted for Immune Evasion. Journal of Virology 91, e00698–00617; 10.1128/JVI.00698-17 (2017).

23 Mutso, M. et al. Reverse genetic system, genetically stable reporter viruses and packaged subgenomic replicon based on a Brazilian Zika virus isolate. Journal of General Virology 98, 2712–2724; 10.1099/jgv.0.000938 (2017).

24 Gabriel, E. et al. Recent Zika Virus Isolates Induce Premature Differentiation of Neural Progenitors in Human Brain Organoids. Cell Stem Cell 20, 397–406; 10.1016/j.stem.2016.12.005 (2017).

25 Retallack, H. a. Zika virus cell tropism in the developing human brain and inhibition by azithromycin. Proceedings of the National Academy of Sciences of the United States of America 113, 14408–14413; 10.1073/pnas.1618029113 (2016).

26 Tang, H. et al. Zika Virus Infects Human Cortical Neural Progenitors and Attenuates Their Growth. Cell Stem Cell 18, 587–590; 10.1016/j.stem.2016.02.016 (2016).

27 Li, C. et al. Zika Virus Disrupts Neural Progenitor Development and Leads to Microcephaly in Mice. Cell Stem Cell 19, 120–126; 10.1016/j.stem.2016.04.017 (2016).

28 Scaturro, P., Cortese, M., Chatel-Chaix, L., Fischl, W. & Bartenschlager, R. Dengue Virus Non-structural Protein 1 Modulates Infectious Particle Production via Interaction with the Structural Proteins. PLOS Pathogens 11, e1005277; 10.1371/journal.ppat.1005277 (2015).

29 Khadka, S. et al. A physical interaction network of dengue virus and human proteins. Mol Cell Proteomics 10, M111 012187; 10.1074/mcp.M111.012187 (2011).

30 Hirsila, M., Koivunen, P., Gunzler, V., Kivirikko, K. I. & Myllyharju, J. Characterization of the human prolyl 4-hydroxylases that modify the hypoxia-inducible factor. J Biol Chem 278, 30772–30780; 10.1074/jbc.M304982200 (2003).

31 Myllyharju, J. Prolyl 4-hydroxylases, key enzymes in the synthesis of collagens and regulation of the response to hypoxia, and their roles as treatment targets. Annals of Medicine 40, 402–417; 10.1080/07853890801986594 (2008).

32 Mizuno, K., Peyton, D. H., Hayashi, T., Engel, J. & Bchinger, H. P. Effect of the -Gly-3(S)-hydroxyprolyl-4(R)-hydroxyprolyl-tripeptide unit on the stability of collagen model peptides. The FEBS Journal 275, 5830–5840; 10.1111/j.1742-4658.2008.06704.x (2008).

33 Ishikawa, Y., Vranka, J., Wirz, J., Nagata, K. & Bachinger, H. P. The rough endoplasmic reticulum-resident FK506-binding protein FKBP65 is a molecular chaperone that interacts with collagens. J Biol Chem 283, 31584–31590; 10.1074/jbc.M802535200 (2008).

34 DiChiara, A. S. et al. Mapping and Exploring the Collagen-I Proteostasis Network. ACS Chem Biol 11, 1408–1421; 10.1021/acschembio.5b01083 (2016).

35 Ishikawa, Y. & Bachinger, H. P. A molecular ensemble in the rER for procollagen maturation. Biochim Biophys Acta 1833, 2479–2491; 10.1016/j.bbamcr.2013.04.008 (2013).

36 Kiriakidis, S. et al. Complement C1q is hydroxylated by collagen prolyl 4 hydroxylase and is sensitive to off-target inhibition by prolyl hydroxylase domain inhibitors that stabilize hypoxia-inducible factor. Kidney International 92, 900–908; 10.1016/j.kint.2017.03.008 (2017).

37 Hudson, D. M. & Eyre, D. R. Collagen prolyl 3-hydroxylation: a major role for a minor post-translational modification? Connective Tissue Research 54, 245–251; 10.3109/03008207.2013.800867 (2013).

38 Reid, K. B. M. Complete amino acid sequences of the three collagen-like regions present in subcomponent C1q of the first component of human complement. Biochemical Journal 179, 367–371; 10.1042/bj1790367 (1979).

39 Pralow, A. et al. Site-specific N-glycosylation analysis of animal cell culture-derived Zika virus proteins. Sci Rep 11, 5147; 10.1038/s41598-021-84682-z (2021).

40 Xu, X. et al. Contribution of intertwined loop to membrane association revealed by Zika virus full-length NS1 structure. EMBO J 35, 2170–2178; 10.15252/embj.201695290 (2016).

41 Sulser, P. et al. HIF hydroxylase inhibitors decrease cellular oxygen consumption depending on their selectivity. The FASEB Journal 34, 2344–2358; 10.1096/fj.201902240R (2020).

42 Sasaki, T., Majamaa, K. & Uitto, J. Reduction of collagen production in keloid fibroblast cultures by ethyl-3,4-dihydroxybenzoate. Inhibition of prolyl hydroxylase activity as a mechanism of action. Journal of Biological Chemistry 262, 9397–9403; 10.1016/S0021-9258(18)48093-3 (1987).

43 Aviner, R., Li, K. H., Frydman, J. & Andino, R. Cotranslational prolyl hydroxylation is essential for flavivirus biogenesis. Nature 596, 558–564; 10.1038/s41586-021-03851-2 (2021).

44 Vasta, J. D. et al. Selective Inhibition of Collagen Prolyl 4-Hydroxylase in Human Cells. ACS Chemical Biology 11, 193–199; 10.1021/acschembio.5b00749 (2016).

45 Wang, J., Buss, J. L., Chen, G., Ponka, P. & Pantopoulos, K. The prolyl 4-hydroxylase inhibitor ethyl-3,4-dihydroxybenzoate generates effective iron deficiency in cultured cells. FEBS Letters 529, 309–312; 10.1016/S0014-5793(02)03389-6 (2002).

46 Warnecke, C. et al. Activation of the hypoxia-inducible factor-pathway and stimulation of angiogenesis by application of prolyl hydroxylase inhibitors. The FASEB Journal 17, 1186–1188; 10.1096/fj.02-1062fje (2003).

47 Li, B., Takeda, K., Yokoyama, S. & Shibahara, S. A prolyl-hydroxylase inhibitor, ethyl-3,4-dihydroxybenzoate, induces haem oxygenase-1 expression in human cells through a mechanism independent of hypoxia-inducible factor-1alpha. J Biochem 144, 643–654; 10.1093/jb/mvn115 (2008).

48 Frakolaki, E. et al. The Role of Tissue Oxygen Tension in Dengue Virus Replication. Cells 7, 241; 10.3390/cells7120241 (2018).

49 Somnuke, P., Hauhart, R. E., Atkinson, J. P., Diamond, M. S. & Avirutnan, P. N-linked glycosylation of dengue virus NS1 protein modulates secretion, cell-surface expression, hexamer stability, and interactions with human complement. Virology 413, 253–264; 10.1016/j.virol.2011.02.022 (2011).

50 Wang, C. et al. Endocytosis of flavivirus NS1 is required for NS1-mediated endothelial hyperpermeability and is abolished by a single N-glycosylation site mutation. PLOS Pathogens 15, e1007938; 10.1371/journal.ppat.1007938 (2019).

51 Bagdonaite, I. & Wandall, H. H. Global aspects of viral glycosylation. Glycobiology 28, 443–467; 10.1093/glycob/cwy021 (2018).

52 Bräutigam, J., Scheidig, A. J. & Egge-Jacobsen, W. Mass spectrometric analysis of hepatitis C viral envelope protein E2 reveals extended microheterogeneity of mucin-type O-linked glycosylation. Glycobiology 23, 453–474; 10.1093/glycob/cws171 (2013).

53 Dechtawewat, T. et al. Mass spectrometric analysis of host cell proteins interacting with dengue virus nonstructural protein 1 in dengue virus-infected HepG2 cells. Biochim Biophys Acta 1864, 1270–1280; 10.1016/j.bbapap.2016.04.008 (2016).

54 Cervantes-Salazar, M. et al. Dengue virus NS1 protein interacts with the ribosomal protein RPL18: this interaction is required for viral translation and replication in Huh-7 cells. Virology 484, 113–126; 10.1016/j.virol.2015.05.017 (2015).

55 Annunen, P., Koivunen, P. & Kivirikko, K. I. Cloning of the alpha subunit of prolyl 4-hydroxylase from Drosophila and expression and characterization of the corresponding enzyme tetramer with some unique properties. J Biol Chem 274, 6790–6796; 10.1074/jbc.274.10.6790 (1999).

56 Ishikawa, Y. et al. Evidence for the major role of PH4αEFB in the prolyl 4-hydroxylation of Drosophila collagen IV. Matrix Biol 141, 101–113; 10.1016/j.matbio.2025.09.002 (2025).

57 Frega, M. et al. Rapid Neuronal Differentiation of Induced Pluripotent Stem Cells for Measuring Network Activity on Micro-electrode Arrays. J Vis Exp 10.3791/54900 (2017).

58 Beers, J. et al. Passaging and colony expansion of human pluripotent stem cells by enzyme-free dissociation in chemically defined culture conditions. Nature Protocols 7, 2029–2040; 10.1038/nprot.2012.130 (2012).

59 Bezemer, B., van Cleef, K. W. R., Overheul, G. J., Miesen, P. & van Rij, R. P. The calcium channel inhibitor lacidipine inhibits Zika virus replication in neural progenitor cells. Antiviral Research 202, 105313; 10.1016/j.antiviral.2022.105313 (2022).

60 Reed, L. J. & Muench, H. A simple method of estimating fifty per cent endpoints. American Journal of Epidemiology 27, 493–497; 10.1093/oxfordjournals.aje.a118408 (1938).

61 Betting, V. et al. A piRNA-lncRNA regulatory network initiates responder and trailer piRNA formation during mosquito embryonic development. RNA 27, 1155–1172; 10.1261/rna.078876.121 (2021).

62 Slonchak, A. et al. Zika virus noncoding RNA suppresses apoptosis and is required for virus transmission by mosquitoes. Nature Communications 11, 2205; 10.1038/s41467-020-16086-y (2020).

63 Livak, K. J. & Schmittgen, T. D. Analysis of Relative Gene Expression Data Using Real-Time Quantitative PCR and the 2(™Delta Delta C(T)) Method. Methods 25, 402–408; 10.1006/meth.2001.1262 (2001).

64 Cox, J. & Mann, M. MaxQuant enables high peptide identification rates, individualized p.p.b.-range mass accuracies and proteome-wide protein quantification. Nature Biotechnology 26, 1367–1372; 10.1038/nbt.1511 (2008).

65 Tyanova, S. et al. The Perseus computational platform for comprehensive analysis of (prote)omics data. Nature Methods 13, 731–740; 10.1038/nmeth.3901 (2016).

66 Szklarczyk, D. et al. The STRING database in 2021: customizable protein–protein networks, and functional characterization of user-uploaded gene/measurement sets. Nucleic Acids Research 49, D605–D612; 10.1093/nar/gkaa1074 (2021).

67 Shannon, P. et al. Cytoscape: A Software Environment for Integrated Models of Biomolecular Interaction Networks. Genome Research 13, 2498–2504; 10.1101/gr.1239303 (2003).

68 Schindelin, J. et al. Fiji: An open-source platform for biological-image analysis. Nature Methods 9, 676–682; 10.1038/nmeth.2019 (2012).

69 Perez-Riverol, Y. et al. The PRIDE database resources in 2022: A hub for mass spectrometry-based proteomics evidences. Nucleic Acids Research 50, D543–D552; 10.1093/nar/gkab1038 (2022).

