## Supplementary Figures 1-7; Supplementary Table captions for "Prolyl hydroxylation is required for maturation and secretion of Zika virus non-structural protein 1"

**Contains:**

- Supplementary figures 1–7
- Supplementary Table captions

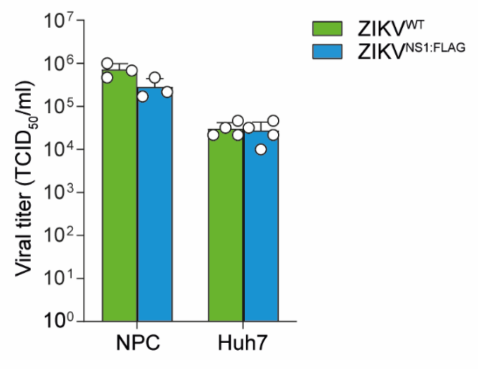


**Supplementary figure 1. Viral titers in samples used for mass spectrometry.**
Infectious titers in the culture supernatant of NPCs and Huh7 cells at 48 hours post-infection with $ZIKV^{WT}$ or $ZIKV^{NS1:FLAG}$ at an MOI of 1.0. Data were log-transformed prior to statistical testing using student's *t*-tests. No significant differences were observed in either cell line. Bars represent the mean and SD of three biological replicates in a single experiment. Individual data points are shown as circles.

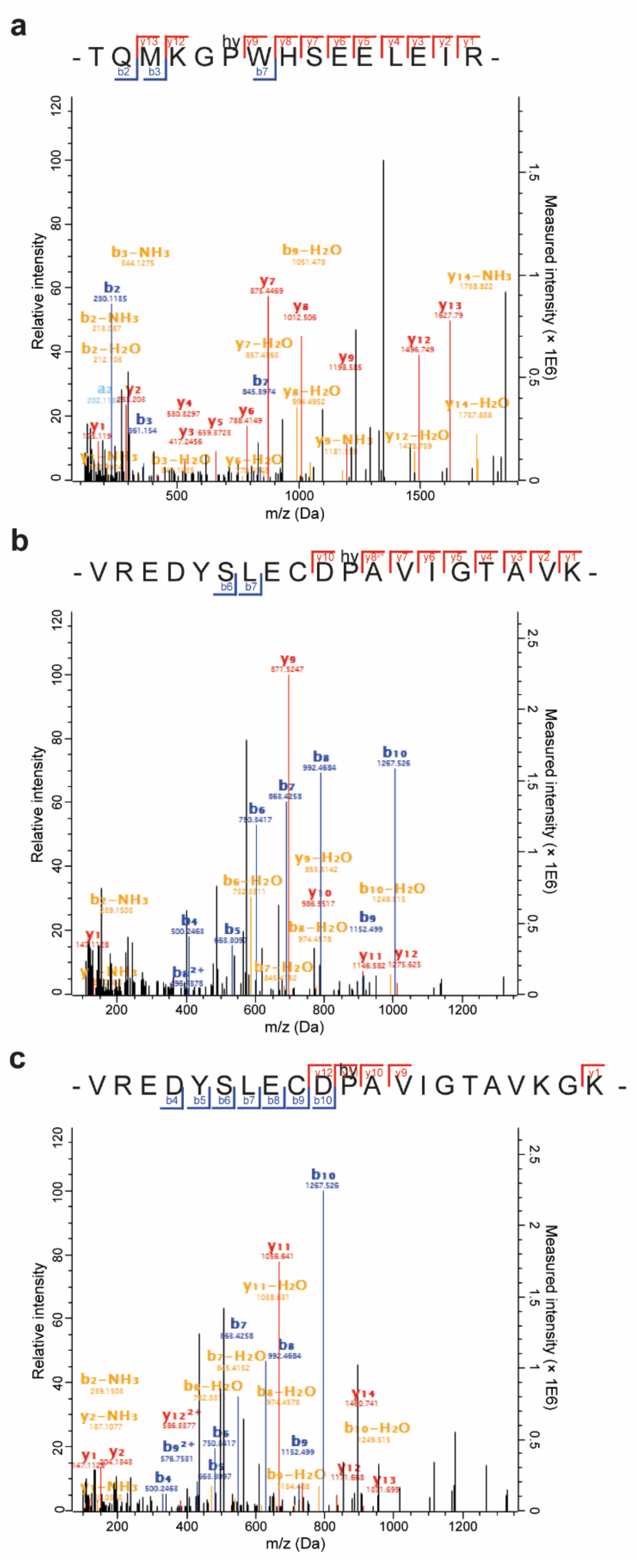


**Supplementary figure 2. Potential hydroxyproline residues in NS1 fragmentation spectra.**
**a–c** Assigned fragmentation spectra for NS1-hydroxyP267 in FLAG IP from ZIKV$^{NS1:FLAG}$ infected NPCs (**a**), NS1-hydroxyP181 (best identified) (**b**), NS1-hydroxyP181 (best localized) (**c**) in FLAG IP from ZIKV$^{NS1:FLAG}$ infected Huh7 cells. The X-axis shows m/z (Da), the left Y-axis indicates the relative intensity, and the right Y-axis shows the measured intensity (× 1E6). Colors indicate the type of ion: red corresponds to a y-ion, blue to a b-ion, and yellow indicates peptide fragment ion with a neutral loss. Identified ions are indicated in the peptide sequence.

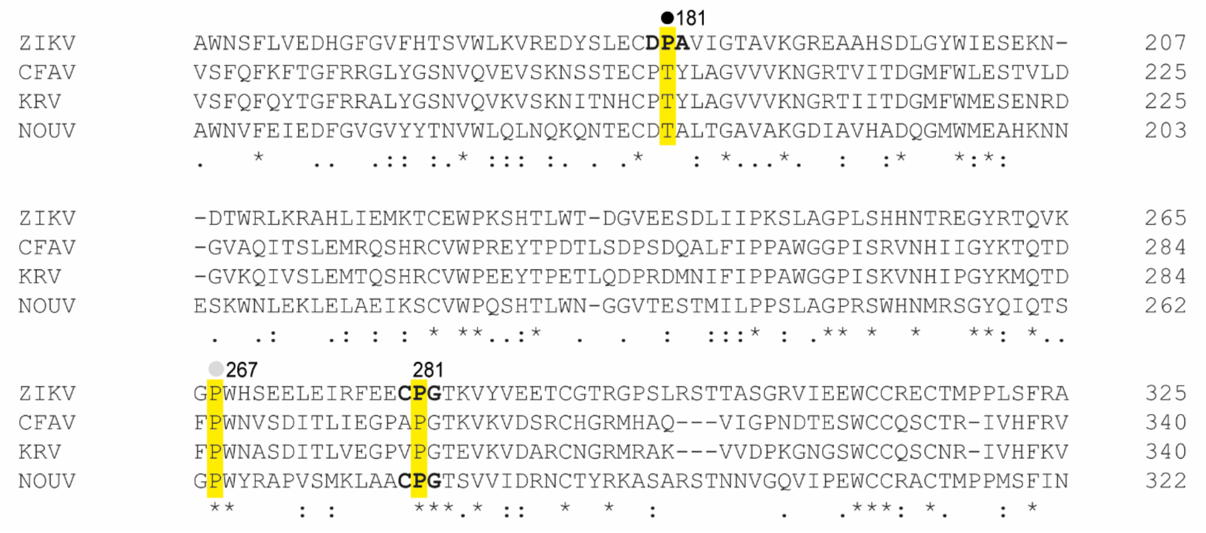


**Supplementary figure 3. Sequence alignment of NS1 from ZIKV and three insect-specific flaviviruses.**

Sequence alignment of amino acids 149–325 of NS1 from ZIKV (NC_012532) with the corresponding sequence of cell-fusing agent virus (CFAV, NC_001564), Kamiti River virus (KRV, NC_005064) and nounané virus (NOUV, NC_033715). Fully conserved residues are indicated by asterisks, conservation of strongly similar amino acids is indicated by colons and conservation of weakly similar amino acids is indicated by dots. A hydroxyproline site in ZIKV NS1 with mass spectrometry support in multiple replicates is indicated with a closed circle and a site with mass spectrometry support in one of three replicates is indicated with a grey circle. Canonical recognition sequences for collagen prolyl 4-hydroxylases are shown in bold. Proline residues selected for further investigation are highlighted in yellow.

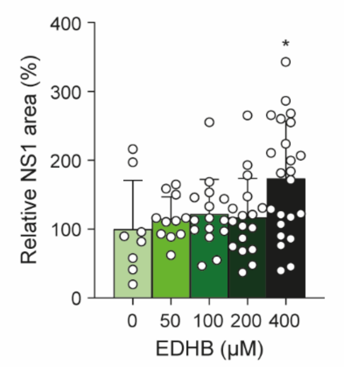


**Supplementary figure 4. NS1-positive cellular area in EDHB-treated cells.**

Signal quantification of the NS1 staining area in Fig. 4a, normalized to the DMSO control for each condition. Data were analyzed with one-way ANOVA with Dunnett's post-hoc test (* $p < 0.05$; ** $p < 0.01$). Bars represent the mean and SD of 39–58 positive cells per condition. Data from individual slides are shown as circles.

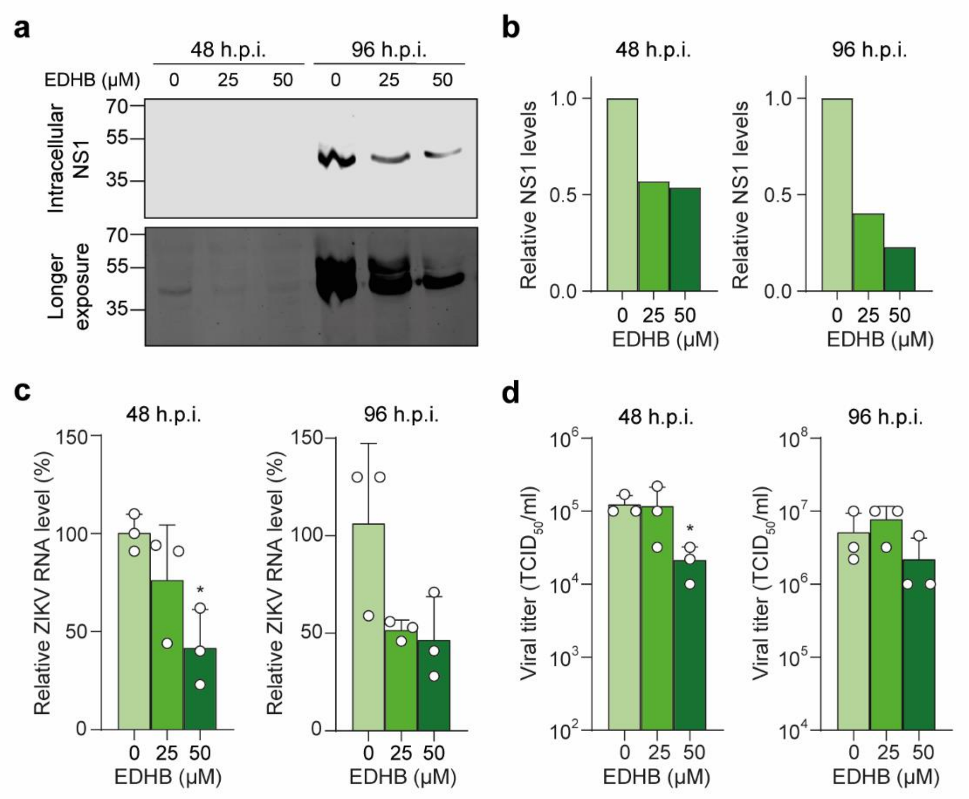


**Supplementary figure 5. Prolonged reduction in ZIKV replication by prolyl hydroxylase inhibition.** **a–d** Intracellular NS1 protein levels (**a, b**), ZIKV RNA levels (**c**) and infectious titers in the culture supernatant of HeLa cells (**d**) at 48 or 96 hours post-infection upon EDHB treatment. Cells were infected with ZIKV$^{NS1:FLAG}$ at an MOI of 0.01 and treated with 25 or 50 µM EDHB or DMSO from 2 hours post-infection onwards. Intracellular NS1 protein was detected using a FLAG antibody (**a**), quantified and normalized to DMSO treated cells (**b**). Bars and dots in panels **c** and **d** represent mean and SD and individual data of three replicate wells, with dots representing the individual data points. Data were analyzed with one-way ANOVA with Dunnett's post-hoc test, using log-transformed data for panel **d** (* $p < 0.05$).

**Linker fragment sequence**

GTGTGGGGAAATCCATGGGTCTGGGTCGGCATGGCATCTCCACCTCCTCGCGGTCCGACCTGGGCATCCGAAGGAGGA
CGTCGTCCACTCGGATGGCTAAGGGAGAGCCACTTTTCTCTCGATTCTCTATCGGAATCTAGGGAGCTCGGATCCAGA
CATGATAAGATACATTGATGAGTTTGGACAAACCACAACTAGAATGCAGTGAAAAAAATGCTTTATTTGTGAAATTTG
TGATGCTATTGCTTTATTTGTAACCATTATAAGCTGCAATAAACAAGTTAACAACAACAATTGCTCGAGGGGGGGCCC
GGTACCTTGAAGCTGTCCCTGATGGTCGTCATCTACCTGCCTGGACAGCATGGCCTGCAACGCGGGCATCCCGATGCC
GCCGGAAGCGAGAAGAATCATAATGGGGAAGGCCATCCAGCCTCGCGTCGGCGCTTAAGACATTGATTATTGACTAGT
TATTAATAGTAATCAATTACGGGGTCATTAGTTCATAGCCCATATATGGAGTTCCGCGTTACATAACTTACGGTAAAT
GGCCCGCCTGGCTGACCGCCCAACGACCCCCGCCCATTGACGTCAATAATGACGTATGTTCCCATAGTAACGCCAATA
GGGACTTTCCATTGACGTCAATGGGTGGAGTATTTACGGTAAACTGCCCACTTGGCAGTACATCAAGTGTATCATATG
CCAAGTCCGCCCCCTATTGACGTCAATGACGGTAAATGGCCCGCCTGGCATTATGCCCAGTACATGACCTTACGGGAC
TTTCCTACTTGGCAGTACATCTACGTATTAGTCATCGCTATTACCATGGTGATGCGGTTTTGGCAGTACACCAATGGG
CGTGGATAGCGGTTTGACTCACGGGGATTTCCAAGTCTCCACCCCATTGACGTCAATGGGAGTTTGTTTTGGCACCAA
AATCAACGGGACTTTCCAAAATGTCGTAACAACTGCGATCGCCCGCCCCGTTGACGCAAATGGGCGGTAGGCGTGTAC
GGTGGGAGGTCTATATAAGCAGAGCTCGTTTAGTGAACCGAGTTGTTGATCTGTGTGAATCA

**Legend**

- Last 22 nucleotides of the ZIKV 3’UTR
- Hepatitis delta virus ribozyme site
- Simian virus 40 late polyadenylation signal
- Cytomegalovirus promoter
- First 22 nucleotides of the ZIKV 5’UTR

**Supplementary figure 6. Linker fragment sequence and annotation.**

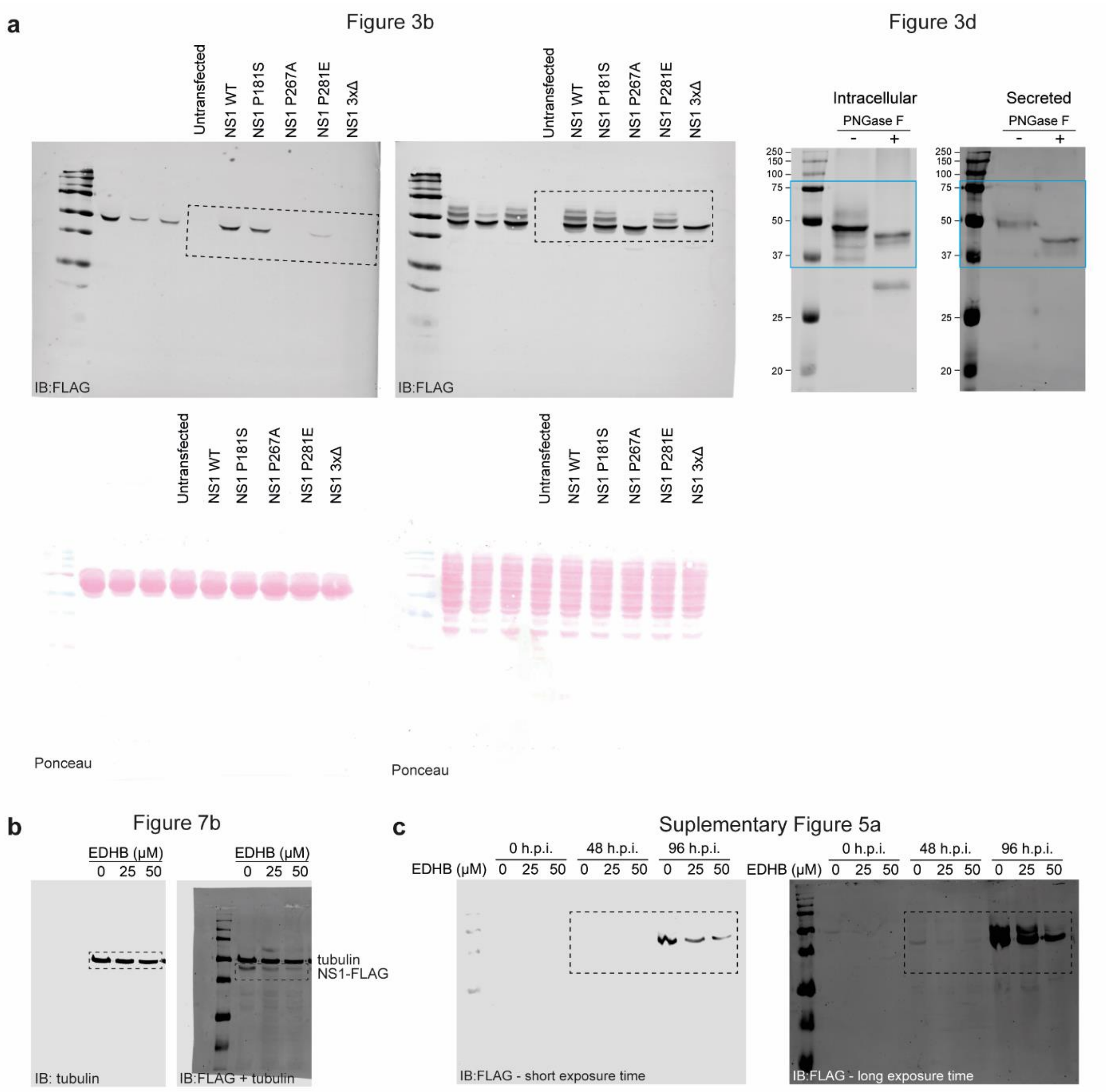


**Supplementary figure 7. Uncropped western blots and Ponceau stains.**
**a** Uncropped western blot images (top) and associated Ponceau stains (bottom) of Fig. 3b (left panels) and Fig. 3d (right panels). **b–c** Uncropped western blot images of Fig. 6b and Supplementary Fig. 5a. Dashed lines indicate the cropped areas.

### Supplementary Table captions

**Supplementary table 1. Oligonucleotides used in this study.**

**Supplementary table 2. Significant NS1 interactors in ZIKV infected NPCs and Huh7 cells.**
The file contains three tabs: one tab shows the NS1 interactors in NPCs and Huh7 cells meeting the fold-change (FC) and *p*-value thresholds (-log10(*p*-value) > 1.3 and log2(FC) > 2 for NPCs and log2(FC) > 3 for Huh cell, respectively). Red font are viral proteins including the NS1 bait; Green font are NS1-interactors shared between NPCs and Huh7 cells. The complete list of proteins identified by mass spectrometry is provided in Supplementary table S3 and S4. The second tab compares the significant NS1 interactors to proteins identified in proteomic studies of dengue virus and ZIKV virus. The third tab summarizes the experimental setup of these studies.

**Supplementary table 3. Quantitative mass spectrometry data of ZIKV infected NPCs.**
Label-free quantitative mass spectrometry data obtained from FLAG IP of lysates from NPCs infected with $ZIKV^{NS1:FLAG}$ or $ZIKV^{WT}$. Columns A–F are the $\log_2$-transformed LFQ intensities. UniProt identifiers or viral protein names, host protein names and gene names are indicated in columns U, W and X, respectively. The $-\log_{10}$ (*p*-value) values from the *t*-tests can be found in column S and a “+” in column J indicates significant outliers. Column T contains the $\log_2$ (fold change) values.

**Supplementary table 4. Quantitative mass spectrometry data of ZIKV infected Huh7 cells.**
Label-free quantitative mass spectrometry data obtained from FLAG IP of lysates from Huh7 cells infected with $ZIKV^{NS1:FLAG}$ or $ZIKV^{WT}$. Columns A-F are the $\log_2$-transformed LFQ intensities. UniProt identifiers or viral protein names, are indicated in column U. The $-\log_{10}$ (*p*-value) values from the *t*-tests can be found in column S and a “+” in column J indicates significant outliers. Column T contains the $\log_2$ (fold change) values.

**Supplementary table 5. Enrichment of Gene Ontology terms in mass spectrometry data of ZIKV infected NPCs.**
Columns C–F contain the enrichment statistics: *Observed gene count*, the number of NS1-interacting proteins associated with the indicated GO term; *Background gene count*, total number of proteins assigned to the GO term; *Strength*, $\log_{10}$ (observed/expected); *False discovery rate*, *p*-values corrected for multiple testing with the Benjamini-Hochberg procedure.

**Supplementary table 6. Enrichment of Gene Ontology terms in mass spectrometry data of ZIKV infected Huh7 cells.**
Columns C–F contain the enrichment statistics: *Observed gene count*, the number of NS1-interacting proteins associated with the indicated GO term; *Background gene count*, total number of proteins assigned to the GO term; *Strength*, $\log_{10}$ (observed/expected); *False discovery rate*, *p*-values corrected for multiple testing with the Benjamini-Hochberg procedure.

**Supplementary table 7. Identification of hydroxyproline sites.**
The file contains 2 tabs with output tables from the MaxQuant search for modified residues; one for Huh7 cells and one for NPCs. Protein and gene names can be found in columns E and F, respectively. Column B indicates which residue of the protein was modified. Column M contains the score for the localization, where a higher score indicates a better identified modified peptide. Localization prob indicates the certainty with which the modification is localized on the peptide, in each of the samples, which is summarized in column AR, where modified sites with their probability are shown in the context of the amino acid sequence.
